# Cardiomyocyte caveolae govern myocardial function and sex-dependent regulation of ventricular compliance and resilience via cavin-1

**DOI:** 10.64898/2026.04.17.717104

**Authors:** BT Quick, HY Khoo, T Bishop, JS Russell, S Niogret, JE Outhwaite, UY Ho, LJ Griffiths, Z Lu, J Rae, NJ Palpant, RG Parton, WG Thomas, JP Headrick, ME Reichelt

**Author notes:** **Corresponding author:** Dr Melissa Reichelt, School of Biomedical Sciences, The University of Queensland, St Lucia, QLD 4072, AUSTRALIA. denotes equal author contributions (BT Quick, HY Khoo, T Bishop).

## Abstract

**Aims:** Caveolae are plasmalemmal microdomains regulating stretch-dependent, nitric oxide (NO), and other signalling pathways governing myocardial structure, function and resilience. We have reported that global deletion of the scaffold protein cavin-1 disrupts caveolar biogenesis and impairs ventricular compliance and tolerance to ischaemic injury. However, cardiomyocyte-specific and sex-dependent roles of cavin-1 and caveolar complexes remain unresolved.

**Methods and Results:** We generated a floxed *Cavin-1* transgenic mouse, enabling cardiomyocyte-specific knockdown via adeno-associated virus (AAV) mediated expression of iCre recombinase driven by a cardiac-specific troponin T promoter. Knockdown was confirmed by RNA, protein, and immunofluorescence analyses, and cardiac function was assessed via echocardiography, left ventricular pressure-volume (PV) catheterisation, and *ex vivo* PV analysis of perfused hearts. Conditionally deleted hearts and myocytes exhibited up to 50% knockdown of *Cavin-1* mRNA together with 15% deficiency in muscle-specific *Caveolin-3*, 70% depletion of caveolae, and mislocalisation of NO synthase (NOS) within cardiomyocytes. This was associated with elevated heart rate and shortened PR interval; reduced intraventricular and systolic blood pressures and peripheral resistance; and sex-dependent impairment of ventricular filling (females only). Diastolic dysfunction was detectable *ex vivo*, to a greater extent in male vs. female hearts. Mechanisms were sex-dependent, linked to interstitial fibrosis in females and NOS overactivity (inhibited by 100 µM L-NAME) in males. Female hearts also exhibited increased susceptibility to ischaemia-reperfusion injury. Coronary function appeared preserved in both sexes, with intact reactive hyperaemic responses.

**Conclusion:** This model identifies cardiomyocyte caveolae and cavin-1 as key determinants of myocardial function and compliance, involving sex-dependent remodelling and NOS signalling. By linking cardiomyocyte disruption to whole-organ and -body dysfunction, this model provides mechanistic insight into impaired function in heart failure and ageing.

**Graphical Abstract:** 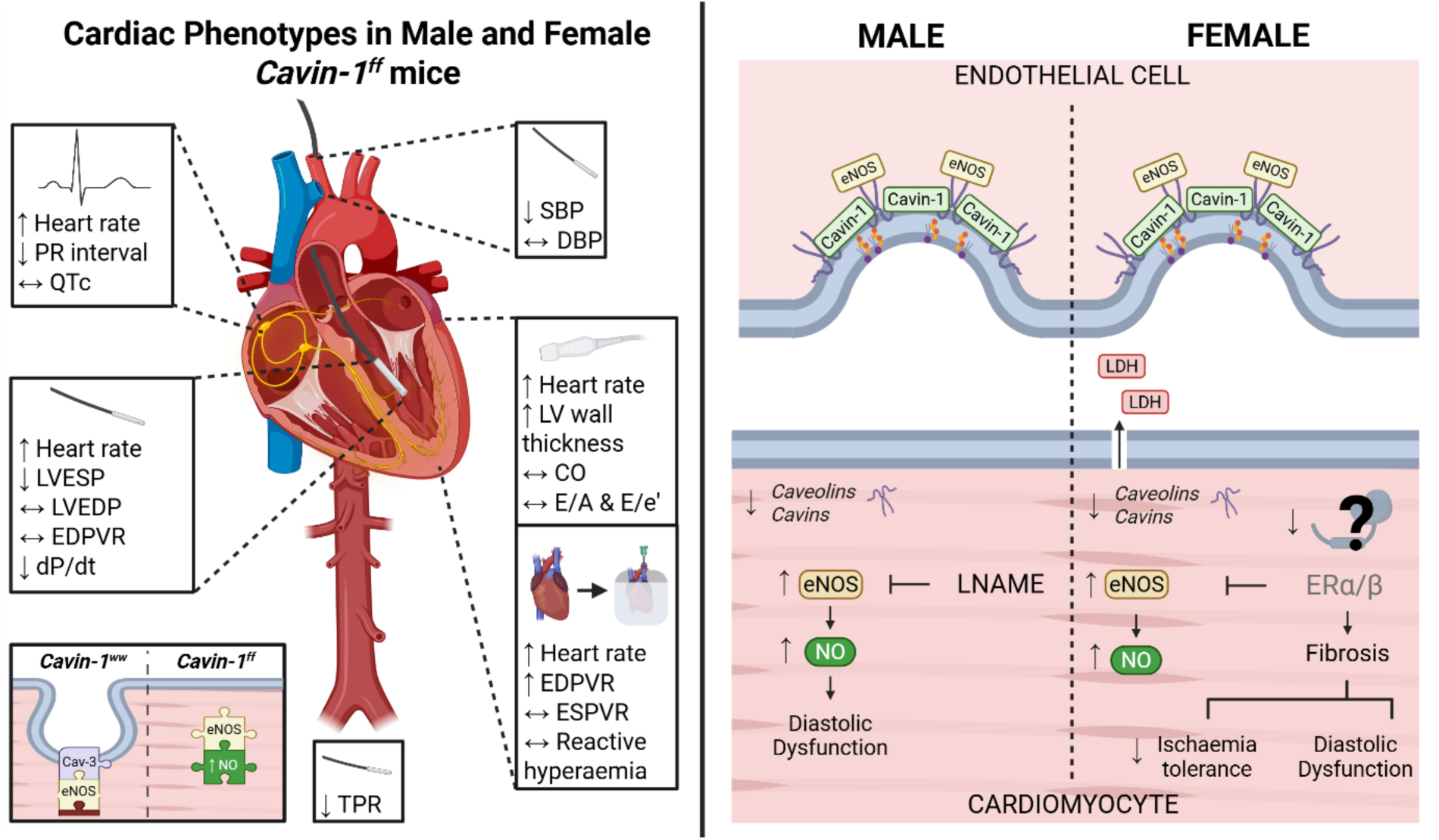

## Introduction

Diastolic dysfunction describes the chronic, progressive stiffening of the heart, functionally manifesting as prolonged relaxation, increased filling pressures, and an inadequate cardiac output despite preserved systolic performance (1). As a hallmark of heart failure with preserved ejection fraction (HFpEF) and cardiac ageing, diastolic dysfunction underscores the significance of ventricular compliance in maintaining cardiovascular health (1, 2). Structural remodelling processes, including fibrosis, which increases myocardial stiffness through excessive extracellular matrix deposition, and cardiac hypertrophy, which reduces chamber compliance by thickening the ventricular wall, further intensify diastolic impairment and accelerate functional decline (3). While specific pathways governing such deterioration remain elusive, diastolic dysfunction has been linked to abnormal formation and function of caveolae – specialised, flask-shaped invaginations of the plasma membrane with multifaceted roles within cardiac cells (4, 5).

The molecular architecture of caveolae is demarcated by two major protein families: caveolins and cavins. Caveolins form oligomeric complexes in the Golgi complex and then these are transported to the plasma membrane where they generate caveolae in cooperation with cavin proteins, particularly cavin1 (6, 7). Studies utilising global knockout mouse models of the ubiquitous caveolin-1 and muscle-specific caveolin-3 highlight their possible roles in regulating cell signalling, endocytosis, and lipid handling, while also serving as inhibitors of nitric oxide synthases (NOS) in the heart (8–12). Consistent with this, loss of caveolin-1 or caveolin-3 severely depletes cardiac caveolae and disrupts membrane receptor and ion channel clustering, alongside a progressive cardiomyopathy evidenced by contractile dysfunction, arrhythmogenesis, and other fatal pathologies (13–16).

Cavins are essential for the recruitment and stabilisation of caveolins and caveolae (17). Functional studies demonstrate that cavin-1 (or polymerase I and transcript release factor, PTRF) is indispensable for caveolar biogenesis, with complete absence of caveolae following *Cavin-1* knockout in mice (18). Cavin-1-dependent cardioprotection arises from its essential role in preserving the spatial fidelity of caveolae-based signalling networks, including β-adrenergic modulation (19), metabolic flux (20), mechanotransduction (21), and ion-channel regulation within cardiomyocytes (22, 23). Caveolae also serve as key organising sites for nitric oxide (NO) signalling, wherein cavin-1 stabilises caveolins that normally restrain NOS activity and prevent excessive NO production (24). Consequently, human mutations in *CAVIN-1* cause congenital generalised lipodystrophy type 4, associated with cardiomyopathy and fatal arrhythmias (25, 26). Similarly, *Cavin-1* null mice exhibit profound cardiovascular abnormalities, including uni/bilateral ventricular hypertrophy, cardiac fibrosis, diastolic dysfunction, impaired vasomotion and pulmonary vascular remodelling, however these phenotypes are notoriously heterogenous across studies (27–29). Crucially, cavin and caveolin proteins in the heart diminish with age, associated with progressive functional decline (30, 31), whereas overexpression of cardiomyocyte-specific caveolin-3 attenuates hypertrophy and post-ischaemic dysfunction (32–34). This suggests that age-dependent deterioration of caveolae formation drives malformations resulting in cardiac stiffening in ageing hearts, predisposing to HFpEF.

Sex differences closely intersect with HF pathophysiology yet remain underexplored at the molecular level. Women exhibit a higher prevalence of HFpEF, accounting for nearly two-thirds of cases, and display distinct patterns of myocardial remodelling and diastolic dysfunction compared with men (35, 36). Sex differences in treatment outcomes are likewise pervasive, with females often experiencing disparities in management and underrepresentation in clinical trials, whereas males paradoxically exhibit higher mortality following diagnosis (37, 38). While the molecular architecture and signalling capacity of caveolae are increasingly recognised as dynamic and hormone-sensitive, how sex-specific factors shape caveolar function in cardiomyocytes, thereby influencing remodelling and stress responses, remains unresolved.

Elucidating the cell type–specific roles of caveolae and individual caveolar proteins in the heart has been limited by the use of global knockout models, in which embryonic deletion yields complex systemic sequelae including metabolic, adipocyte, pulmonary, and immune dysfunction (together with potential developmental adaptations) that preclude interrogation of cardiomyocyte influences (18, 39). Here, employing postnatal adeno-associated virus (AAV)-mediated cavin-1 knockdown as a means to selectively perturb caveolar architecture in cardiomyocytes, we examined how caveolae influence haemodynamic function, diastolic compliance, and ischaemia tolerance, with attention to sex-specific differences in these phenotypes.

## Methods

### Mice

This work was undertaken with approval from the University of Queensland Animal Ethics Committee, under project numbers 2021/AE000567 and 2023/AE000402. The *Cavin-1* transgenic mouse model was developed in collaboration with the Queensland Facility of Advanced Genome Editing (QFAGE) using CRISPR gene-editing. Four single guide RNAs and a repair template were designed, which includes the LoxP sequence either side of the exon inserted in place of the endogenous exon 1 of *Cavin-1*. These components were then delivered to ∼200 CD1 ova via microinjection, and male founders were backcrossed with CD1 females to generate *Cavin-1* flox/flox (*Cavin-1*^ff^) mice. Genotyping primers can be found in the supplement (Table S1). Male and female *Cavin-1* homozygous wild-type (*Cavin-1*^ww^) and *Cavin-1*^ff^ littermates were used for all experiments.

### AAV constructs

The adeno-associated virus (AAV) construct was designed in collaboration with Vector Biolabs (Malvern, PA, USA) to co-express a codon-improved Cre recombinase (iCre) and enhanced green fluorescent protein (eGFP) driven by a cardiac troponin T (cTnT) promoter. The cTnT promoter enables cardiomyocyte-specific gene expression, with a short 2A peptide (T2A) permitting co-translational production of separate proteins from the single transcript. Expression of the construct was stabilised by the woodchuck hepatitis post-transcriptional regulatory element (WPRE). The construct (pAAV-cTnT-iCre-T2A-eGFP-WPRE) was utilised with plasmids for pAAV2/9n (rep2/cap9; #V005569, NovoPro) and pHelper (Addgene) for virus production.

### AAV production

Virus preparations were from the University of Queensland’s Viral Vector Core facility, as per the protocol of Kimura et al. (40). Briefly, 70-90% confluent AAV-293 cells were transfected with pAAV-cTnT-iCre-eGFP, pAAV2/9n and pHelper. Viruses were harvested and purified using PEG precipitation, chloroform extraction, two-phase separation in aqueous solution, and discontinuous gradients of iodixanol. Purified AAV were then concentrated using an Amicon^TM^ 100 kDa filter (Merck Millipore). Titre was initially quantified by reverse transcription quantitative polymerase chain reaction (RT-qPCR) of *Cre* primers and later by digital droplet PCR. Aliquoted AAVs were stored at −80°C until use and kept on ice while defrosted for administration.

### Neonatal virus administration

One day after birth (P1), neonatal mice were anaesthetised in ice for 60 sec, and 30 μL of sterile diluted virus (2×10^11^ vgc/mouse; 1X PBS) was injected into the temporal vein. Mice recovered under a heat lamp until mobile and were rolled in soiled bedding prior to being returned to their mother to minimise cannibalisation.

### Conscious assessment of ECG activity

Electrocardiographic recordings were obtained from conscious, unrestrained mice using the ECGenie Clinic system (Mouse Specifics Inc., Framingham, MA, USA). Mice were acclimated to the recording platform prior to data acquisition to minimise stress-related variability. Electrocardiographic signals were captured via footplate electrodes embedded in the recording chamber, allowing non-invasive detection of cardiac electrical activity through the paws. Recordings were performed at a sampling rate of 2 kHz, enabling high-resolution capture of murine ECG intervals. Data were collected over a 15-min period per animal, and segments free of motion artifacts were selected for analysis. Parameters including heart rate, PR interval, QRS duration, and QT interval were extracted.

### Echocardiography

Cardiac function was assessed *in vivo* in 14-15 week-old mice with a Vevo 3100 Imaging System (VisualSonics, Toronto, Canada), using the 55-MHz (MX550D) transducer. Anaesthesia was induced with 4% isoflurane in O_2_ (0.8 L/min) and maintained at 1.5-2%. Heart rate (HR), respiration, and electrocardiography were recorded, with body temperature preserved at 37.0±0.5°C using a heat pad and lamp, as required. Two-dimensional B-mode images were acquired in the parasternal long-axis view and used to derive left ventricular ejection fraction (LVEF), end-systolic volume (LVESV), end-diastolic volume (LVEDV), and cardiac output (normalised to body weight; CI). Global longitudinal strain was also calculated from long-axis images using Vevo Strain speckle-tracking analysis software. M-mode echocardiography in the parasternal short-axis view was used to obtain LV wall thickness. An apical four-chamber view was utilised to identify the mitral valve for assessment of both pulse wave Doppler, measuring blood flow during early ventricular filling (E) and atrial contraction (A), and tissue Doppler imaging, capturing mitral annular motion during early diastolic filling (e’) and atrial contraction (a’). Recorded images were analysed using Vevo LAB 3.1.1 software (VisualSonics, Toronto, Canada) in a blinded fashion. All parameters were measured for at least 3 cardiac cycles, with averaged values taken for analysis.

### Tissue collection

Mice were humanely euthanised either by cervical dislocation or following injection with ketamine/xylazine (50 and 20 mg/mL, respectively; refer to timeline in Figure 1A). Tissues for histology and immunofluorescence analyses were fixed in 4% PFA overnight and transferred to 70% ethanol prior to embedding. Hearts used for molecular analysis were snap frozen in liquid N_2_ and stored at −80°C until processed.

**Figure 1:**
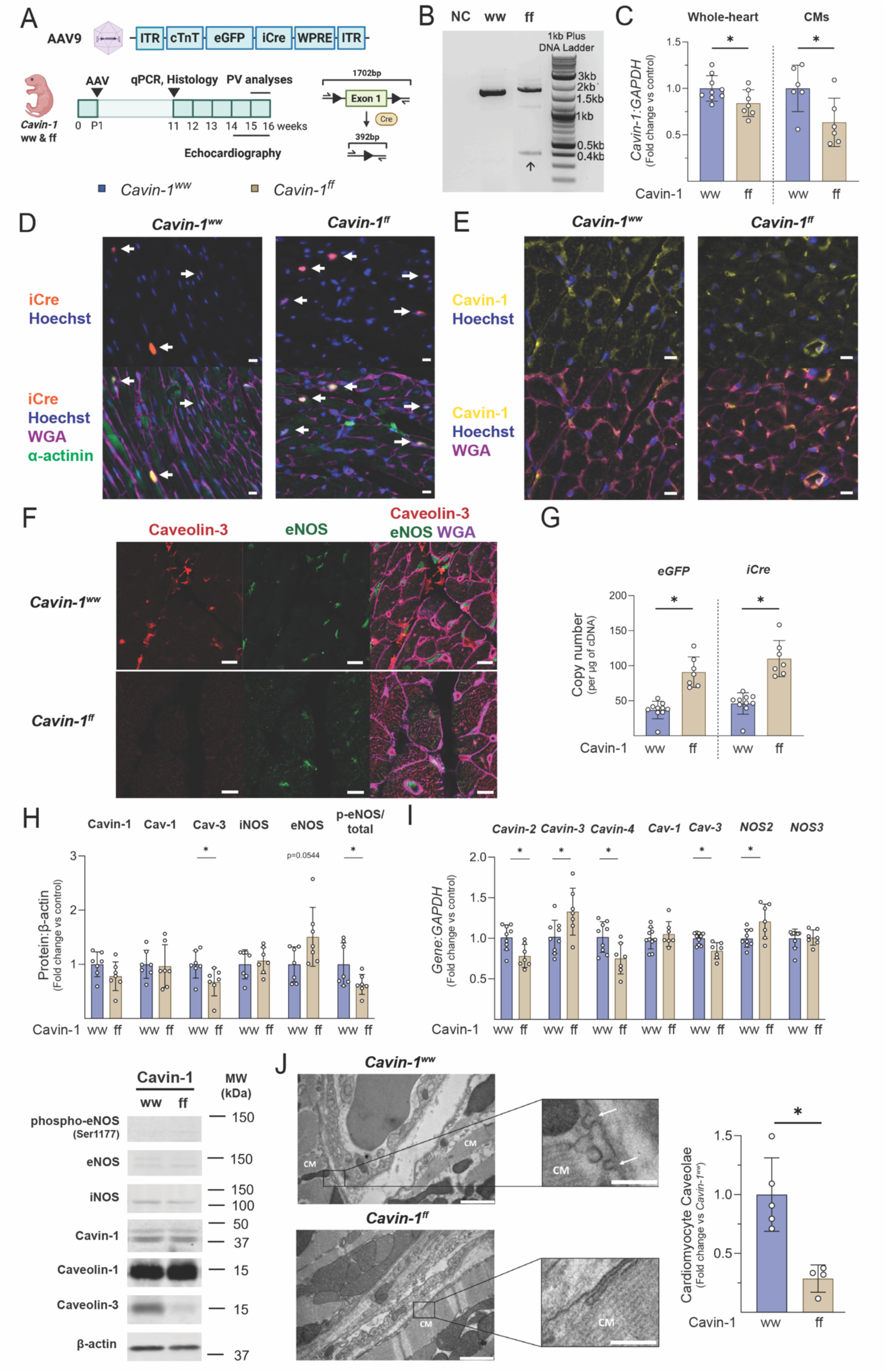
AAV-inducible Cre recombinase drives partial cardiomyocyte cavin-1 knockdown and caveolae disruption, with data pooled for males and females. (A) Schematic illustrating the timeline of neonatal AAV-mediated cavin-1 knockdown and subsequent functional and molecular analyses. (B) Agarose gel electrophoresis demonstrating Cre-mediated DNA recombination after deletion of *Cavin-1* exon 1, as indicated by the arrow. (C) Quantitative PCR analysis of *Cavin-1* mRNA expression in whole-heart tissue (*n*_ww_=9, *n*_ff_=7) and cardiomyocyte-enriched fractions (*n*_ww_=6, *n*_ff_=6), normalised to *GAPDH*. (D, E) Confocal micrographs showing expression of Cre recombinase (orange, D) and cavin-1 (yellow, E), co-stained with wheat germ agglutinin (WGA, purple), α-actinin (green), and Hoechst (blue). Scale bars: 10μm. (F) Immunofluorescent staining of caveolin-3 (red), endothelial nitric oxide synthase (eNOS; green), and WGA (purple) in paraffin mouse heart sections. Scale bars: 10 μm. (G) mRNA expression of exogenous *iCre* and *eGFP* transgenes, quantified as copy number per μg of cDNA (*n*_ww_=10, *n*_ff_=7). (H) Immunoblot analysis of whole-heart protein expression of cavin-1, caveolin-1 (Cav-1), caveolin-3 (Cav-3), inducible NOS (iNOS), and eNOS, normalised to β-actin. Phosphorylated eNOS (ser1177) is shown relative to total eNOS. Blots are provided in the Supplement Figure S2 (*n*_ww_=7, *n*_ff_=7). (I) Whole-heart mRNA expression of *Cavin-2, Cavin-3, Cavin-4, Cav-1, Cav-3, NOS2*, and *NOS3* (*n*_ww_=9, *n*_ff_=7). (J) Representative transmission electron micrographs of *Cavin-1*^ww^ and *Cavin-1*^ff^ hearts. Scale bars: 1000 nm (main panels), 250 nm (insets). White arrows indicate caveolae at the cardiomyocyte plasma membrane. Data are presented as mean ± SD and were analysed by unpaired two-tailed t-tests. Sexes are pooled, with sex-stratified data show in in Table S1. *=*p*<0.05, individual comparisons as indicated. Abbreviations: CM=cardiomyocytes, Cav-1/3=Caveolin-1/3, WGA=wheat germ agglutinin.

### Pressure-volume *in vivo* analyses

A subset (*n*=4-6) of 14-16 week-old *Cavin-1*^ww^ and *Cavin-1*^ff^ mice were anaesthetised with 4% isoflurane in 0.8-1 mL/min O_2_, prior to intubation. Mice were positioned supine, with body temperature maintained at 37.0±0.5°C (monitored via rectal probe). For pressure assessment, a Millar Pressure-Volume catheter (ADI Instruments, SPR-839) was advanced into the left ventricle via the right carotid artery and aortic valve, secured with three 5-0 Prolene suture anchoring loops (Ethicon, Somerville, NJ, USA). Baseline measurements were recorded, followed by occlusions achieved by pinching the ventilator tube for 3-5 sec at end-expiration, transiently decreasing venous return and ventricular preload to allow determination of load-independent indices (performed in triplicate). Data acquisition and analysis were performed in LabChart v8.1.

### Langendorff isolated heart model

Directly following *in vivo* catheterisation, hearts were isolated and immediately submerged in ice-cold modified Krebs-Henseleit buffer, and aortas cannulated on a blunted 21-gauge needle and retrograde perfused (coronary pressure of 80 mmHg) on a Langendorff apparatus, as detailed previously (29). Coronary flow was measured via an in-line Doppler flow probe (Transonic Systems Inc., Clifton, NJ, USA) and left-ventricular pressures were recorded using an intraventricular balloon connected to a pressure transducer. Data was recorded on a PowerLab system (ADInstruments Pty Ltd.; Bella Vista, Australia), with ventricular pressures digitally processed to provide peak rates of pressure development (dP/dt_max_) and relaxation (dP/dt_min_).

#### Left-ventricular pressure-volume relationships

Ventricular end-diastolic and end-systolic pressure-volume relationships (EDPVR, ESPVR) were assessed *ex vivo* to identify intrinsic cardiac changes free from extrinsic influences, as described previously (29). Briefly, following cannulation and equilibration, the ventricular balloon was briefly inflated to a systolic pressure of ∼100 mmHg, and then deflated to 0 mmHg and stabilised for 10 min. After a 5 min equilibration period, hearts were treated with 100 µM of the NOS inhibitor N-Ω-Nitro-L-arginine methyl ester hydrochloride (L-NAME, Sigma Aldrich; St. Louis, MO, USA) or vehicle (buffer). The 100μM concentration is widely used to achieve robust NOS inhibition in *ex vivo* cardiovascular preparations (29, 41–43). Hearts were then paced 8 min into equilibration at a mean rate of 470 beats/min, to normalise rate between wild-type and floxed groups (floxed mice exhibiting elevated heart rate). Baseline efflux of LDH was assessed in cardiac effluent collected during equilibration (stored at −80°C). After stabilisation at 0 mmHg systolic pressure, balloon volume was incrementally increased in 2.65 µL steps using a 500 µL threaded syringe (Hamilton Co; Reno, NV, USA), with function assessed after 2 min at each volume. Experiments were terminated when end-diastolic pressure exceeded 20 mmHg, or after 15 increments (whichever occurred first).

#### Reactive Hyperaemia

For assessment of coronary function, reactive hyperaemic responses were assessed following 15 and 30 sec occlusions. Flow repayment, peak flow and time-to-peak were quantified, and total hyperaemic response was calculated as the area under the flow curve (AUC) over a 2 min period.

#### Ischaemia-reperfusion injury

A separate cohort of perfused hearts were allowed to stabilise for 20 min before electrical pacing at 470 bpm. After an additional 10 min stabilisation period, global ischaemia was induced under normothermic conditions for 25 min. This was succeeded by 45 min reperfusion with oxygenated buffer. Coronary effluent was collected at three time points: 2 min prior to onset of ischaemia, and at 2 and 45 min into the reperfusion phase.

Analysis of all Langendorff data was performed blinded.

### LDH efflux and troponin assay

Lactate dehydrogenase activity was quantified in cardiac effluent collected from perfused hearts using the CytoTox 96® Non-Radioactive Cytotoxicity Assay (ProMega) following manufacturer’s instructions. A standard curve was generated using L-LDH from bovine heart (Sigma-Aldrich; L2625).

Myocardial release of cardiac Troponin I (cTnI) was assessed via enzyme-linked immunosorbent assay (ELISA, LifeDiagnostics, Inc.; CTNI-1-US). Coronary effluent samples were thawed and assayed for cTnI according to manufacturer instructions. Absorbance was determined spectrophotometrically at 450 nM and efflux normalised to coronary flow rate and heart weight.

### Cardiomyocyte purification

Adult cardiomyocytes (week 11) were isolated as described previously (44), with slight modification. Briefly, mice were anaesthetised via a nose cone with 4% isoflurane. The chest cavity was opened, descending aorta and inferior vena cava cut, and EDTA buffer (7 mL) was injected into the right ventricle for 1 min. The aorta was then clamped and EDTA buffer (10 mL), perfusion buffer (3 mL) and pre-warmed collagenase buffer (37°C, 40 mL) were subsequently perfused into the left ventricle at a rate of 1 mL/min. The clamp was removed, and ventricular tissue teased into pieces in 3 mL collagenase buffer and gently triturated for 2 min. The enzymatic digestion was terminated by addition of 5 mL stop solution. The cell suspension was filtered through a 200 µM strainer (Invitrogen, SKU – 43-50200-03) and underwent 3 x 20 min of gravity settling in perfusion buffer. The supernatant (non-myocyte fraction) was collected, the pellet (myocyte fraction) resuspended in perfusion buffer, and both samples centrifuged at 300*g* for 15 min. Supernatants were discarded and pellets again resuspended in perfusion buffer. A 400 μL volume of Trizol^TM^ reagent was added to both fractions to await molecular analyses.

### qPCR

Total RNA was extracted from hearts and digested myocytes using TRIzol reagent (ThermoFisher, Australia), treated with TURBO DNase™ (ThermoFisher), and reverse-transcribed into cDNA using the High-Capacity cDNA Reverse Transcription Kit (Applied Biosystems). RT-qPCR was performed with Fast SYBR^TM^ Green Master Mix (ThermoFisher) and analysed by a QuantStudio^TM^ 6 Flex Real-Time PCR System. Primer sequences can be found in the Supplement (Table S2). Exogenous *iCre* recombinase and *eGFP* transgene expression were quantified by a standard curve generated from a plasmid of known concentration followed by log transformation. Endogenous expression was quantified by the ΔΔ-Ct method with *GAPDH* as a housekeeping gene. Results were taken as null if multi-peak melt curves were observed.

### Western blotting

Tissues were homogenised (IKA T-10 basic homogeniser) in buffer containing 100 mM Tris-HCl (pH 7.0) and 5 mM each of EGTA and EDTA, supplemented with protease inhibitors (cOmplete, Roche; Basel, Switzerland) and phosphatase inhibitors (PhosSTOP^TM^, Roche; Basel, Switzerland). Protein concentration was measured via colourimetric BCA assay (BioRad; #500013, #500014). Extracts were denatured in Laemmli buffer (95°C, 5 min), before separation on a 12% acrylamide gel. BioRad Precision Plus Protein^TM^ Dual Colour Standards was used as a molecular weight marker. Proteins were transferred to Immobilon-FL polyvinylidene fluoride membranes (Merck Millipore) using a tank-based wet transfer electrophoretic system. Membranes were blocked for 1 hr at room temperature in a 1:1 solution of Intercept blocking buffer (LI-COR) and PBS, and incubated overnight at 4°C with primary antibodies in blocking buffer: anti-Ptrf1/Cavin-1 (Rabbit, Sigma-Aldrich ABT131, 1:1000), anti-caveolin-3 (Mouse, BDBiosciences 610421, 1:10000), anti-caveolin-1 (Rabbit, ThermoFisher PA5-17447, 1:1000), anti-iNOS (Rabbit, ThermoFisher PA1-036, 1:1000), anti-eNOS (Rabbit, ThermoFisher PA3-031A, 1:1000), anti-phospho-eNOS (Ser1177; Rabbit, Cell Signaling 9571S, 1:1000), and anti-β-actin (Mouse, ThermoFisher MA1-140, 1:10000). A series of 5×10 min room temperature washes with 0.1% PBS-T preceded the addition of secondary antibodies diluted 1:10,000 in Intercept/PBS (LI-COR; IRDye 800CW Donkey anti-rabbit 926-32213, IRDye 680RD Donkey anti-mouse 926-68072), and incubation for 1 hr at room temperature away from light. Blots were imaged with an Odyssesy CLx Scanner (LI-COR). Densitometric analysis was performed on the combined intensity of the doublets corresponding to eNOS and cavin-1, likely post-transational modifications, representing total protein abundance.

### Histology and immunofluorescence

Fixed tissues were dehydrated in serially decreasing concentrations of ethanol and embedded in paraffin wax. Sections of 5 μm were obtained and underwent Massons Trichrome staining for fibrosis. In brief, deparaffinised and rehydrated sections were stained with Weigert’s haematoxylin (10 min) and Biebrich scarlet-acid fuchsin (15 min), differentiated in phosphomolybdic-phosphotungstic acid (15 min), then transferred to aniline blue (3 min). Brightfield images were acquired at 40X magnification using a Zeiss Axioscan Z1 slide scanner. Fibrosis analysis was semi-automated using the colour deconvolution plugin on ImageJ in a blinded fashion, as described previously (45). For immunofluorescence, deparaffinised and rehydrated sections were subjected to antigen retrieval at 110°C for 20 min in Tris-EDTA buffer pH 9.0. Non-specific staining was blocked using 10% goat serum (2 hrs, room temperature; Jackson ImmunoResearch) prior to incubation with primary antibodies: anti-Ptrf1/Cavin-1 (Rabbit, Sigma-Aldrich ABT131, 1:200), anti-iCre (Cell Signaling Technology D7L7L, 1:100), anti-α-actinin (Mouse, Sigma-Aldrich, 1:200), anti-eNOS (Rabbit, ThermoFisher PA3-031A, 1:250), and anti-caveolin-3 (Mouse, BDBiosciences 610421, 1:500) diluted in 2% goat serum, overnight at 4°C. Sections were incubated with secondary antibodies (anti-mouse 488, ThermoFisher A21042, 1:500; anti-rabbit 555, ThermoFisher, 1:500) for 2 hrs at room temperature and mounted in ProLong Diamond (ThermoFisher P36970). Images were acquired at 63x magnification using the Zeiss LSM900 AiryScan 2.

### Electron microscopy

Transmission electron microscopy (TEM) was employed to visualise caveolae. Hearts dedicated to TEM were excised, and dissected into ∼1 mm^3^ pieces, before 2 hr fixation with 2.5% glutaraldehyde in PBS at room temperature. Samples were processed for Epon embedding, as described previously (46).

The number of caveolae at the cardiomyocyte membrane were quantitated from 20 representative sections per heart, imaged at 8000x magnification. All quantitation was performed blind independently by two researchers with coded samples.

### Statistics

All data are expressed as mean ± standard deviation. Comparisons were made using unpaired two-tailed t-tests, one-way ANOVA, two-way ANOVA or repeated measures ANOVA with appropriate post hoc correction, as indicated. Figure 1 data were analysed using unpaired two-tailed t-tests, or Mann-Whitney test for nonparametric data (assessed by Shapiro-Wilk), with sex differences assessed via two-way ANOVA followed by uncorrected Fisher’s least significant difference (LSD) test. For Figure 2, qPCR and heart weight data were evaluated using two-way ANOVA with uncorrected Fisher’s LSD, whereas body weight comparisons employed Tukey’s multiple comparisons test. Pressure–volume and echocardiographic parameters were analysed using two-way ANOVA to assess genotype and sex effects, followed by uncorrected Fisher’s LSD. For *ex vivo* pressure–volume analyses, Langendorff-derived functional metrics were assessed using two-way repeated measures ANOVA with Tukey’s post hoc correction, whereas heart rate and area-under-the-curve (AUC) values were compared using two-way ANOVA with uncorrected Fisher’s LSD. Ischaemia–reperfusion data, including functional and LDH measurements, were analysed using two-way repeated measures ANOVA with Tukey’s post hoc correction. When data were non-normal prior to 2-way ANOVA, values were log₁₀-transformed to satisfy assumptions of normality. Statistical significance was defined as *p*<0.05. All analyses were performed using GraphPad Prism statistical packages (v10; GraphPad Software, Inc.).

**Figure 2:**
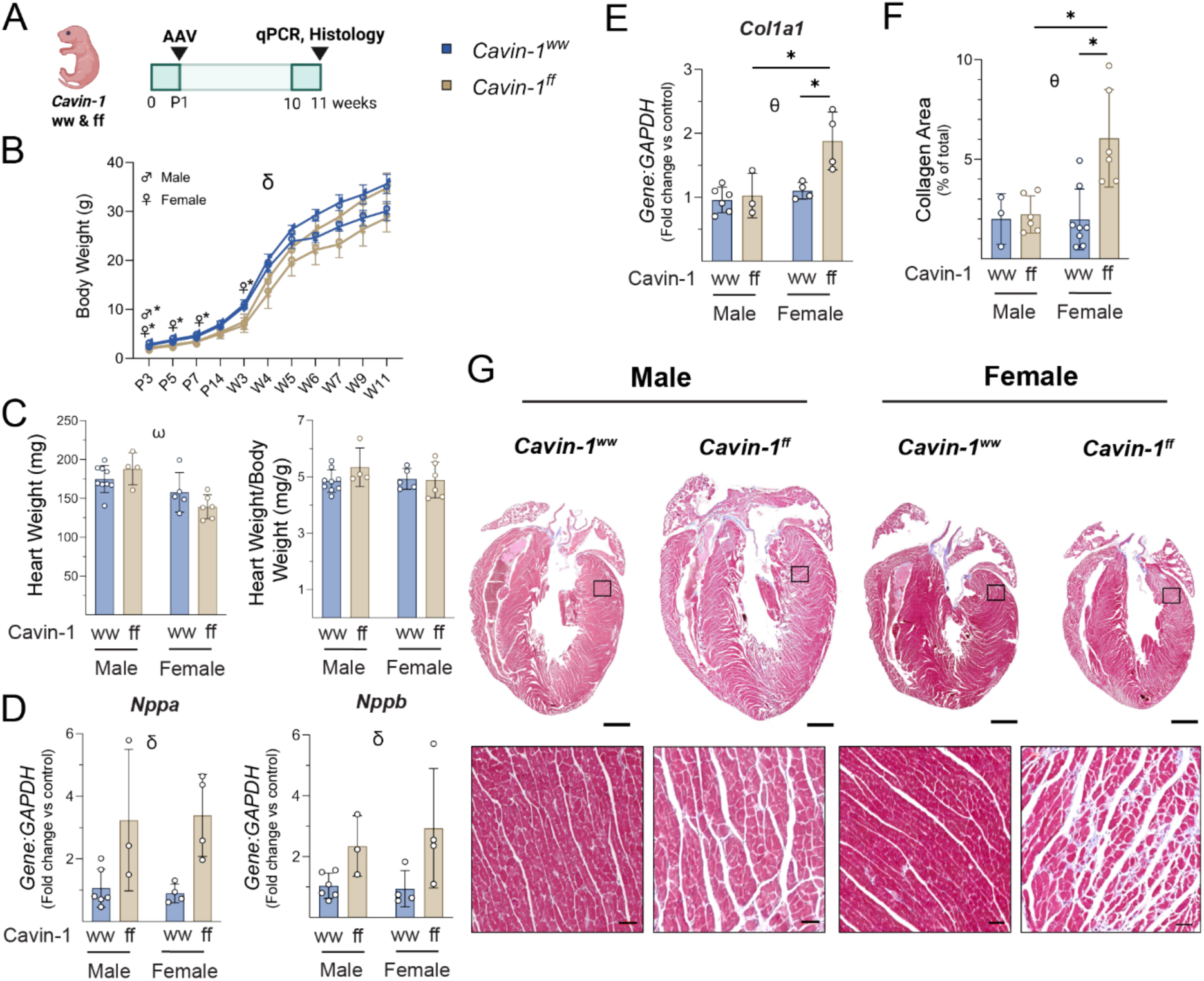
Cavin-1 knockdown in cardiomyocytes elicits female-restricted increases in collagen expression and cardiac fibrosis. (A) Schematic of timelines for analyses. (B) Sex-stratified body weights, (C) raw (left graph) and normalised heart weights (to body weight; right graph). *Cavin-1*^ww^ males, *n*=9; *Cavin-1*^ww^ females, *n*=6; *Cavin-1*^ff^ males, *n*=4; *Cavin-1*^ff^ females, *n*=6. (D) qPCR expression analysis of cardiac stress biomarkers, and (E), (F) fibrotic markers. *Cavin-1*^ww^ males, *n*=6; *Cavin-1*^ww^ females, *n*=3/4; *Cavin-1*^ff^ males, *n*=3; *Cavin-1*^ff^ females, *n*=4. (G) Quantification of collagen via Massons Trichrome staining, and representative longitudinal sections from *Cavin-1*^ww^ and *Cavin-1*^ff^ female and male hearts. Scale bar is 1 mm, scale bar of insets is 50 μm. *Cavin-1*^ww^ males, *n*=3; *Cavin-1*^ww^ females, *n*=8; *Cavin-1*^ff^ males, *n*=6; *Cavin-1*^ff^ females, *n*=6. Data are presented as mean ± SD. qPCR and heart weight data were evaluated using two-way ANOVA with uncorrected Fisher’s LSD, while body weight comparisons employed Tukey’s multiple comparisons test. *, *p*<0.05, individual comparisons as indicated; θ, *p*_interaction_<0.05; δ, *p*_genotype_<0.05; ω, *p*_sex_<0.05.

## Results

### Cardiomyocyte-targeted *Cavin-1* knockdown drives caveolae disruption

To determine whether cardiomyocyte-specific knockdown of cavin-1 disrupts caveolar integrity and NOS signalling, hearts were isolated from *Cavin-1*^ww^ and *Cavin-1*^ff^ mice injected with cardiac-targeted AAV9-iCre (Figure 1A). All data in Figure 1 is presented as sex-pooled data due to lack of sex-specific differences. PCR amplification of genomic DNA confirmed partial exon 1 deletion of *Cavin-1*, generating a 392bp product (Figure 1B). Quantitative PCR analysis indicated a ∼20% reduction of *Cavin-1* mRNA in whole-heart lysates from *Cavin-1*^ff^ mice (*p*=0.0417), which was more evident in enzymatically isolated cardiomyocyte fractions (∼40% knockdown, *p*=0.0327; Figure 1C). Immunofluorescence demonstrated nuclear iCre expression localised within cardiomyocytes across both genotypes (Figure 1D). In wild-type hearts, cavin-1 colocalised with WGA at both cardiomyocyte and endothelial membranes. In *Cavin-1*^ff^ hearts, cavin-1 staining was absent from cardiomyocyte membranes but retained in endothelial cells (Figure 1E), indicating cell-type-specific recombination in cardiomyocytes. We conducted further immunological staining to assess changes in endothelial NOS (eNOS) expression with cavin-1 deficiency. In *Cavin-1*^ww^ hearts, caveolin-3 was detected along the cardiomyocyte sarcolemma, whereas eNOS predominantly localised to endothelial cell clusters, with weaker cardiomyocyte plasma membrane-associated expression (Figure 1F). However, cardiomyocyte cavin-1 deficiency abolished caveolin-3 staining, and eNOS exhibited diffuse cytosolic distribution. Dysregulation of eNOS appears conserved between *Cavin-1*^ff^ males and females, as confirmed by a loss of the striated sarcolemmal signal (Figure S1).

The AAV-mediated delivery of *eGFP* and *iCre* transgenes was robust and consistent between males and females (*eGFP: p*_interaction_=0.9285, *p*_sex_=0.2222; *iCre: p*_interaction_=0.7060, *p*_sex_=0.1404; Table S1), although ∼2-fold elevated in *Cavin-1*^ff^ hearts (*p*=0.0001 for both, Figure 1G).

Densitometric analysis of cavin-1 from whole-heart fractions revealed no significant reduction in protein expression (p=0.1220), likely reflecting masking by preserved endothelial cavin-1. However, cavin-1*-*deficient hearts had significantly lower caveolin-3 protein (∼42%; *p*=0.0262), while caveolin-1 (*p*=0.8504) expression was unchanged, and total eNOS increased (0.5-fold) despite a ∼37% reduction in the ratio of its Ser1177 phosphorylated form (*p*=0.0443; Figure 1H; full blots in Figure S2). *Cavin-1*^ff^ hearts exhibited downregulation of mRNA for *Cavin-2* (∼22%; *p*=0.0089)*, Cavin-4* (∼26%; *p*=0.0154), and *Caveolin-3* (∼15%; *p*=0.0031), alongside up-regulation of *Cavin-3* (0.3-fold; *p*=0.0243), whereas *Caveolin-1* remained unmodified (*p*=0.4571; Figure 1G). *NOS2* (iNOS) mRNA was also elevated (0.2-fold; *p*=0.0230), while *NOS3* (eNOS) was unaffected (*p*=0.8125). Importantly, a 2-way ANOVA identified no significant interaction between sex and genotype for gene or protein expression (Table S3).

Electron microscopy of myocardial tissue confirmed depletion of caveolae at the cardiomyocyte membrane of *Cavin-1*^ff^ hearts, with a ∼70% reduction in relative abundance (*p*=0.0036; Figure 1J). Taken together, we show that partial knockdown of cardiomyocyte cavin-1 is sufficient to impair caveolae formation and dysregulate cardiac NO.

### *Cavin-1* deficiency induces sex-specific cardiac fibrosis

We next investigated the impact of depletion of cardiomyocyte caveolae on body weight and cardiac structure. Cavin-1 knockdown at P1 (Figure 2A) reduced body weight in neonates (Male: P3, *p*=0.0286; Female: P3, *p*=0.0002, P5, *p*=0.0014, P7, *p*=0.0295, week 3, *p*=0.0121; Figure 2B). Heart weights were affected only by sex (*p*_interaction_=0.0681, *p*_sex_=0.0007, *p*_genotype_=0.7316), and this difference was not evident when heart weight was normalised to body weight (*p*_interaction_=0.2356, *p*_genotype_=0.3015, *p*_sex_=0.4017; Figure 2C).

Cavin-1 deficiency also led to sex-independent up-regulation in mRNA expression of cardiac hypertrophy markers *Nppa* (*p*_interaction_=0.7850, *p*_genotype_=0.0015, *p*_sex_=0.9897) and *Nppb* (*p*_interaction_=0.6159, *p*_genotype_=0.0027, *p*_sex_=0.9566; Figure 2D). Expression of fibrosis markers *Col3a1, Acta2, Tgfβ1,* and *PDGFRα* was unchanged (Figure S3). However, *Col1a1* expression was significantly elevated in *Cavin-1*^ff^ females only (*p*_interaction_=0.0303, *p*=0.0024 vs. *Cavin-1*^ww^ females, *p*=0.0021 vs. *Cavin-1*^ff^ males; Figure 2E). Massons Trichrome staining of transverse ventricular sections uncovered severe interstitial fibrosis in the ventricles of *Cavin-1*^ff^ female but not male hearts (*p*_interaction_=0.0190, *p*=0.0003 vs. *Cavin-1*^ww^ females, *p*=0.0009 vs. *Cavin-1*^ff^ males; Figure 2F and 2G). This confirms a crucial role for cardiomyocyte caveolae in supressing fibrotic remodelling in female hearts.

### Echocardiography uncovers divergent sex-specific phenotypes with *Cavin-1* deficiency

To evaluate the functional consequences of cardiomyocyte Cavin 1 depletion, Echocardiographic and conscious electrocardiographic analyses in 14-15 week-old males and females revealed an elevated heart rate with cavin-1 deficiency, associated with a shortened PR interval (males and females) and ST segment (males only), without changes in QRS or QTc durations (Table 1).

**Table 1:**
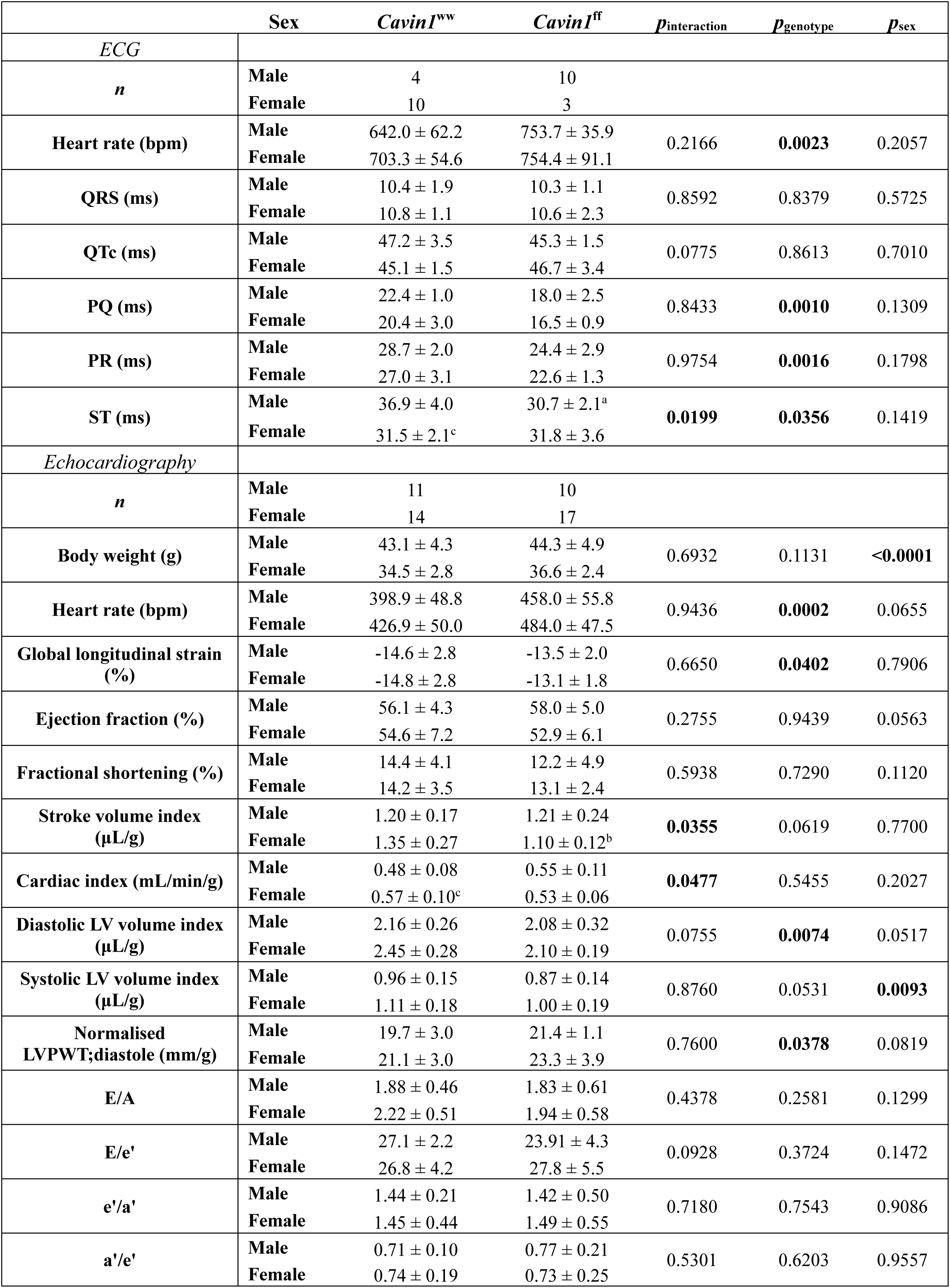
Echocardiographic and conscious electrocardiographic assessments of cardiac function in 14-15 week-old *Cavin-1*^ww^ and *Cavin-1*^ff^ mice. . Data were analysed by 2-way ANOVA with uncorrected Fisher’s LSD for post hoc analyses. Non-normal data was log_10_-transformed and re-analysed. Abbreviations: LV=left ventricle, LVPWT=left ventricular posterior wall thickness. a = *p*<0.05, *Cavin-1*^ff^ males vs *Cavin-1*^ww^ males; b = *p*<0.05, *Cavin-1*^ff^ females vs *Cavin-1*^ww^ females; c = *p*<0.05, *Cavin-1*^ww^ females vs *Cavin-1*^ww^ males.

|  | Sex | *Cavin1*<sup>ww</sup> | *Cavin1*<sup>ff</sup> | *p*<sub>interaction</sub> | *p*<sub>genotype</sub> | *p*<sub>sex</sub> |
| --- | --- | --- | --- | --- | --- | --- |
| *ECG* |  |  |  |  |  |  |
| *n* | Male | 4 | 10 |  |  |  |
|  | Female | 10 | 3 |  |  |  |
| Heart rate (bpm) | Male | 642.0 ± 62.2 | 753.7 ± 35.9 | 0.2166 | **0.0023** | 0.2057 |
|  | Female | 703.3 ± 54.6 | 754.4 ± 91.1 |  |  |  |
| QRS (ms) | Male | 10.4 ± 1.9 | 10.3 ± 1.1 | 0.8592 | 0.8379 | 0.5725 |
|  | Female | 10.8 ± 1.1 | 10.6 ± 2.3 |  |  |  |
| QTc (ms) | Male | 47.2 ± 3.5 | 45.3 ± 1.5 | 0.0775 | 0.8613 | 0.7010 |
|  | Female | 45.1 ± 1.5 | 46.7 ± 3.4 |  |  |  |
| PQ (ms) | Male | 22.4 ± 1.0 | 18.0 ± 2.5 | 0.8433 | **0.0010** | 0.1309 |
|  | Female | 20.4 ± 3.0 | 16.5 ± 0.9 |  |  |  |
| PR (ms) | Male | 28.7 ± 2.0 | 24.4 ± 2.9 | 0.9754 | **0.0016** | 0.1798 |
|  | Female | 27.0 ± 3.1 | 22.6 ± 1.3 |  |  |  |
| ST (ms) | Male | 36.9 ± 4.0 | 30.7 ± 2.1<sup>a</sup> | **0.0199** | **0.0356** | 0.1419 |
|  | Female | 31.5 ± 2.1<sup>c</sup> | 31.8 ± 3.6 |  |  |  |
| *Echocardiography* |  |  |  |  |  |  |
| *n* | Male | 11 | 10 |  |  |  |
|  | Female | 14 | 17 |  |  |  |
| Body weight (g) | Male | 43.1 ± 4.3 | 44.3 ± 4.9 | 0.6932 | 0.1131 | **<0.0001** |
|  | Female | 34.5 ± 2.8 | 36.6 ± 2.4 |  |  |  |
| Heart rate (bpm) | Male | 398.9 ± 48.8 | 458.0 ± 55.8 | 0.9436 | **0.0002** | 0.0655 |
|  | Female | 426.9 ± 50.0 | 484.0 ± 47.5 |  |  |  |
| Global longitudinal strain (%) | Male | -14.6 ± 2.8 | -13.5 ± 2.0 | 0.6650 | **0.0402** | 0.7906 |
|  | Female | -14.8 ± 2.8 | -13.1 ± 1.8 |  |  |  |
| Ejection fraction (%) | Male | 56.1 ± 4.3 | 58.0 ± 5.0 | 0.2755 | 0.9439 | 0.0563 |
|  | Female | 54.6 ± 7.2 | 52.9 ± 6.1 |  |  |  |
| Fractional shortening (%) | Male | 14.4 ± 4.1 | 12.2 ± 4.9 | 0.5938 | 0.7290 | 0.1120 |
|  | Female | 14.2 ± 3.5 | 13.1 ± 2.4 |  |  |  |
| Stroke volume index (μL/g) | Male | 1.20 ± 0.17 | 1.21 ± 0.24 | **0.0355** | 0.0619 | 0.7700 |
|  | Female | 1.35 ± 0.27 | 1.10 ± 0.12<sup>b</sup> |  |  |  |
| Cardiac index (mL/min/g) | Male | 0.48 ± 0.08 | 0.55 ± 0.11 | **0.0477** | 0.5455 | 0.2027 |
|  | Female | 0.57 ± 0.10<sup>c</sup> | 0.53 ± 0.06 |  |  |  |
| Diastolic LV volume index (μL/g) | Male | 2.16 ± 0.26 | 2.08 ± 0.32 | 0.0755 | **0.0074** | 0.0517 |
|  | Female | 2.45 ± 0.28 | 2.10 ± 0.19 |  |  |  |
| Systolic LV volume index (μL/g) | Male | 0.96 ± 0.15 | 0.87 ± 0.14 | 0.8760 | 0.0531 | **0.0093** |
|  | Female | 1.11 ± 0.18 | 1.00 ± 0.19 |  |  |  |
| Normalised LVPWT;diastole (mm/g) | Male | 19.7 ± 3.0 | 21.4 ± 1.1 | 0.7600 | **0.0378** | 0.0819 |
|  | Female | 21.1 ± 3.0 | 23.3 ± 3.9 |  |  |  |
| E/A | Male | 1.88 ± 0.46 | 1.83 ± 0.61 | 0.4378 | 0.2581 | 0.1299 |
|  | Female | 2.22 ± 0.51 | 1.94 ± 0.58 |  |  |  |
| E/e' | Male | 27.1 ± 2.2 | 23.91 ± 4.3 | 0.0928 | 0.3724 | 0.1472 |
|  | Female | 26.8 ± 4.2 | 27.8 ± 5.5 |  |  |  |
| e'/a' | Male | 1.44 ± 0.21 | 1.42 ± 0.50 | 0.7180 | 0.7543 | 0.9086 |
|  | Female | 1.45 ± 0.44 | 1.49 ± 0.55 |  |  |  |
| a'/e' | Male | 0.71 ± 0.10 | 0.77 ± 0.21 | 0.5301 | 0.6203 | 0.9557 |
|  | Female | 0.74 ± 0.19 | 0.73 ± 0.25 |  |  |  |

Global longitudinal strain was modestly but significantly reduced following cavin-1 knockdown, indicating impaired myocardial deformation (Table 1). However, left ventricular ejection fraction and fractional shortening remained unchanged, despite a *Cavin-1*^ff^ female-restricted reduction in stroke volume indices (*p*=0.002 vs. *Cavin-1*^ww^ females; Table 1). Elevated heart rate, alongside variable stroke volume, contributed to trending increases in normalised cardiac output in *Cavin-1*^ff^ males (*p*=0.094 vs. *Cavin-1^ww^* males) but not females (*p*=0.2668 vs. *Cavin-1*^ww^ females; Table 1). End-diastolic volume was significantly reduced with cavin-1 knockdown, independent of sex. In contrast, end-systolic volume index was higher in females, while there was a modest, non-significant trend toward impairment in *Cavin-1*^ff^ *mice* (*p*_sex_=0.0531; Table 1). There was also significant left ventricular posterior wall hypertrophy with cavin-1 deficiency in both sexes, while no diastolic function parameters were altered.

### Pressure-volume analyses reveal left ventricular underloading with preserved diastolic function

*In vivo* ventricular catheterisation identified marked alterations in systemic and cardiac haemodynamics (Table 2). *Cavin-1*^ff^ mice again exhibited increased heart rate under anaesthesia, with significantly reduced systolic, but preserved diastolic blood pressure. This resulted in lower mean arterial pressure, indicating (in the absence of changes in ventricular ejection) a 30-35% reduction in peripheral vascular resistance. Parallel to these systemic effects, left ventricular end-systolic pressure declined by 13-15mmHg, resulting in a significant reduction in developed pressure despite unchanged end-diastolic pressure.

**Table 2:** In vivo pressure–volume loop analysis of cardiac function in 14-16 week-old *Cavin-1*^ww^ and *Cavin-1*^ff^ mice. Data were analysed by 2-way ANOVA with uncorrected Fisher’s LSD for post hoc analyses. Abbreviations: LV=left ventricle, EDPVR=end-diastolic pressure-volume relationship, PRSW=preload-recruitable stroke work, ESPVR=end-systolic pressure-volume relationship.

|  | Sex | *Cavin1<sup>ww</sup>* | *Cavin1<sup>ff</sup>* | *p*<sub>interaction</sub> | *p*<sub>genotype</sub> | *p*<sub>sex</sub> |
| --- | --- | --- | --- | --- | --- | --- |
| *Load-Dependent* |  |  |  |  |  |  |
| *n* | Male | 4 | 6 |  |  |  |
|  | Female | 5 | 5 |  |  |  |
| Heart Rate (bpm) | Male | 448.3 ± 38.2 | 497.8 ± 52.2 | 0.8509 | **0.0436** | 0.8609 |
|  | Female | 448.6 ± 44.4 | 490.3 ± 9.6 |  |  |  |
| Systolic Blood Pressure (mmHg) | Male | 76.2 ± 9.7 | 66.1 ± 11.0 | 0.7127 | **0.0176** | 0.1705 |
|  | Female | 71.5 ± 10.4 | 57.94 ± 10.1 |  |  |  |
| Diastolic Blood Pressure (mmHg) | Male | 49.4 ± 12.9 | 41.8 ± 11.9 | 0.9980 | 0.1291 | 0.5172 |
|  | Female | 46.2 ± 11.3 | 38.7 ± 7.2 |  |  |  |
| Mean Arterial Pressure (mmHg) | Male | 58.3 ± 11.8 | 49.9 ± 11.5 | 0.9932 | 0.0845 | 0.3718 |
|  | Female | 54.1 ± 11.5 | 45.7 ± 7.6 |  |  |  |
| Total Peripheral Resistance (PRU) | Male | 4.48 ± 2.48 | 3.01 ± 1.00 | 0.9666 | **0.0315** | 0.5594 |
|  | Female | 4.83 ± 0.86 | 3.41 ± 1.34 |  |  |  |
| LV End-Systolic Pressure (mmHg) | Male | 80.4 ± 9.7 | 67.2 ± 10.8 | 0.7622 | **0.0032** | 0.2749 |
|  | Female | 76.9 ± 9.2 | 61.1 ± 6.7 |  |  |  |
| LV End-Diastolic Pressure (mmHg) | Male | 3.94 ± 3.11 | 2.71 ± 1.36 | 0.8279 | 0.2108 | 0.7966 |
|  | Female | 3.97 ± 1.55 | 3.10 ± 0.84 |  |  |  |
| LV Developed Pressure (mmHg) | Male | 76.5 ± 8.3 | 64.5 ± 10.1 | 0.7170 | **0.0038** | 0.2313 |
|  | Female | 73.0 ± 9.8 | 58.0 ± 5.9 |  |  |  |
| dP/dt<sub>max</sub> (mmHg/s) | Male | 7114.3 ± 910.7 | 5431.5 ± 1734.4 | 0.9102 | **0.0318** | 0.0362 |
|  | Female | 5476.6 ± 2084.3 | 3950.2 ± 623.1 |  |  |  |
| dP/dt<sub>min</sub> (mmHg/s) | Male | -7800.3 ± 1274.0 | -5879.2 ± 1822.6 | 0.8150 | **0.0144** | 0.0291 |
|  | Female | -6104.4 ± 1483.8 | -4489.6 ± 741.0 |  |  |  |
| Stroke Work (mmHg*mL) | Male | 2.75 ± 0.82 | 1.48 ± 0.43 | 0.0648 | **0.0153** | 0.2921 |
|  | Female | 1.92 ± 0.33 | 1.72 ± 0.77 |  |  |  |
| E<sub>a</sub> (mmHg/μL) | Male | 1755.0 ± 235.6 | 1775.0 ± 432.8 | 0.1230 | 0.1454 | 0.2974 |
|  | Female | 2308.6 ± 647.1 | 1662.2 ± 354.7 |  |  |  |
| Tau (ms) | Male | 5.65 ± 0.78 | 5.91 ± 0.98 | 0.6793 | 0.8729 | **0.0145** |
|  | Female | 7.03 ± 1.32 | 6.92 ± 0.60 |  |  |  |
| *Load-Independent* |  |  |  |  |  |  |
| *n* | Male | 4 | 5 |  |  |  |
|  | Female | 3 | 5 |  |  |  |
| EDPVR Coefficient (α) | Male | 20.4 ± 4.1 | 30.0 ± 11.5 | 0.2798 | 0.8207 | 0.2392 |
|  | Female | 35.5 ± 19.9 | 29.8 ± 13.9 |  |  |  |
| PRSW (mmHg*mL) | Male | 52.1 ± 13.8 | 50.9 ± 13.8 | 0.9738 | 0.8157 | 0.0595 |
|  | Female | 40.3 ± 5.5 | 38.8 ± 9.79 |  |  |  |
| ESPVR | Male | 1664.1 ± 1068.9 | 1801.3 ± 859.9 | 0.6773 | 0.9545 | 0.1938 |
|  | Female | 1311.5 ± 588.6 | 1130.8 ± 270.7 |  |  |  |

Indices of contractility and relaxation (dP/dt_max_ and dP/dt_min_) were uniformly depressed with cavin-1 knockdown in males and females, consistent with reduced loading conditions (Table 2). Stroke work was also diminished in *Cavin-1*^ff^ mice. E_a_, a measure of arterial elastance, and Tau were not impacted by loss of cavin-1, however Tau was higher in females compared to males. Importantly, when assessed independently of loading conditions, EDPVR, ESPVR, and preload recruitable stroke work were preserved under conditions of caveolar depletion (Table 2). However, EDPVR interpretation is complicated by low intraventricular volumes and pressures in *Cavin-1*^ff^ mice shifting the pressure-volume loop downward and leftward, with diastolic data clustering near the asymptote where pressure changes minimally with volume.

### Diastolic impairment and sex dependency in NO-induced dysfunction *ex vivo*

Hearts were isolated immediately following *in vivo* PV analysis and perfused on a Langendorff apparatus for pre/afterload-independent *ex vivo* assessment of compliance, force development, and reactive hyperaemia (Figure 3A and 3B). At baseline, *Cavin-1*^ff^ hearts exhibited intrinsically elevated heart rate (*p*_genotype_=0.0135; Figure 3C) that was NOS-independent (Figure S4). Baseline LDH efflux (Figure 3D) was largely unmodified, although there was a trend towards greater LDH release in *Cavin-1*^ff^ (*p*_genotype_=0.0661) and female hearts (*p*_sex_=0.0607). Troponin showed a similar trend (Figure S5). *Ex vivo* PV analysis revealed pronounced diastolic dysfunction in *Cavin-1*^ff^ hearts, evidenced by leftward displacement of the EDPVR. This was most apparent in males, particularly during the early filling phase, where the initial EDPVR slope was nearly doubled compared with females (0–15.87 μL: *Cavin-1*^ff^ male = 13.8-fold, *Cavin-1*^ff^ female =2.3-fold; Figure 3E and 3F). In *Cavin-1*^ff^ male hearts, NOS inhibition with L-NAME partially normalised chamber compliance, producing a rightward shift of the EDPVR at lower volumes, although increased stiffness persisted at higher volumes. Consistent with these observations, AUC was elevated confirming diastolic stiffness in *Cavin-1*^ff^ males (8.4-fold vs. *Cavin-1*^ww^, *p*=0.0486), while NOS inhibition significantly reduced the overall burden of diastolic dysfunction (*p*_interaction_=0.0112, *p*=0.0041 vs. *Cavin-1*^ff^).

**Figure 3:**
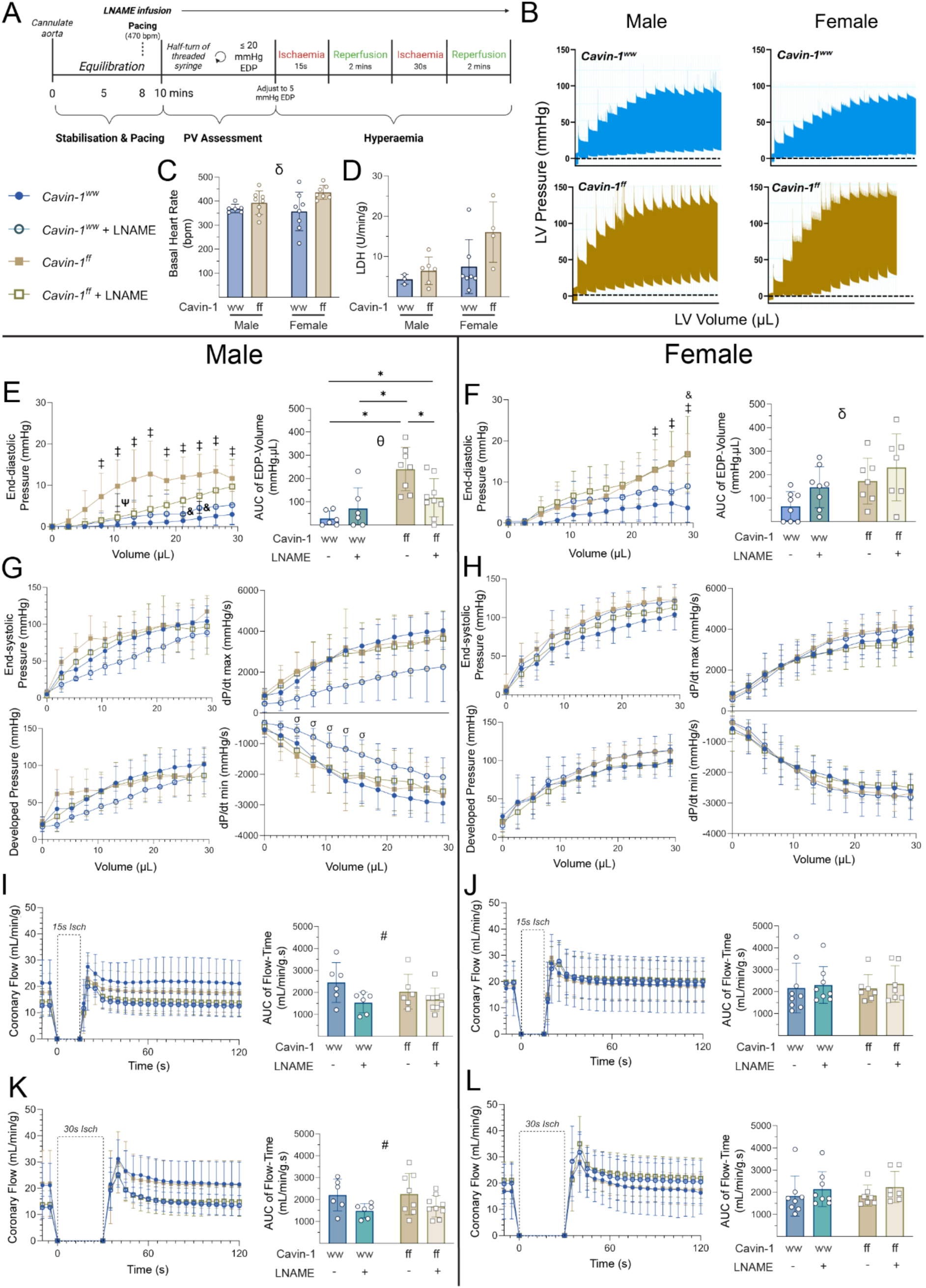
*Ex vivo* Langendorff analysis reveals significant intrinsic diastolic dysfunction in *Cavin-1*^ff^ hearts, with NOS-dependent modulation of compliance and flow restricted to males. (A) Schematic of pressure-volume and reactive hyperaemia protocols in isolated hearts treated with 100 μM L-NAME or vehicle. (B) Representative left ventricular pressure traces from male (left) and female (right) *Cavin-1*^ww^ (top) and *Cavin-1*^ff^ (bottom) hearts. (C) Pre-paced heart rate from vehicle-treated cohorts (*Cavin-1*^ww^ males, *n*=6; *Cavin-1*^ww^ females, *n*=8; *Cavin-1*^ff^ males, *n*=8; *Cavin-1*^ff^ females, *n*=7) and (D) LDH efflux (*Cavin-1*^ww^ males, *n*=3; *Cavin-1*^ww^ females, *n*=7; *Cavin-1*^ff^ males, *n*=6; *Cavin-1*^ff^ females, *n*=4). (E, F) End-diastolic pressure-volume relationship and area-under-the-curve (AUC) analyses of pressure traces in male (left) and female (right) hearts ± L-NAME. Male: *n*_ww_=6, *n*_ww+L-NAME_=6, *n*_ff_=8, *n*_ff+L-NAME_=9. Female: *n*_ww_=8, *n*_ww+L-NAME_=8, *n*_ff_=7, *n*_ff+L-NAME_=7. (G, H) End-systolic, developed pressure, dP/dt_min_ and dP/dt_max_ parameters. (I-L) Reactive hyperaemia responses following 15s (I,J) and 30s (K, l) occlusions in untreated *Cavin-1*^ww^ (*n*_male_=6, *n*_female_=9) and *Cavin-1*^ff^ (*n*_male_=6, *n*_female_=7) and 100 µM L-NAME treated *Cavin-1*^ww^ (*n*_male_=6, *n*_female_=7) and *Cavin-1*^ff^ (*n*_male_=9, *n*_female_=7) hearts. Data are presented as mean ± SD. All data were analysed by 2-way ANOVA, with Tukey’s post hoc correction for functional metrics, while uncorrected Fisher’s LSD was used post hoc for LDH efflux, heart rate, and AUC. *, *p*<0.05, individual comparisons as indicated; θ, *p*_interaction_<0.05; δ, *p*_genotype_<0.05; #, *p*_L-NAME treatment_<0.05. Repeated measures ANOVA, ‡, *p*_ff_<0.05 vs *Cavin-1*^ww^; &, *p*_ff+L-NAME_<0.05 vs *Cavin-1*^ww^; σ, *p*_ww+L-NAME_<0.05 vs *Cavin-1*^ww^; Ψ, *p*_ff+L-NAME_<0.05 vs *Cavin-1*^ff^.

In females, the impact of cardiomyocyte cavin 1 depletion on diastolic phenotype was comparatively modest, with a smaller increase in EDPVR AUC (2.6-fold vs. *Cavin-1*^ww^; *p*_genotype_=0.0145). Importantly, L-NAME treatment did not alter the EDPVR in *Cavin-1*^ff^ females (*p*_interaction_=0.7625, *p*_treatment_=0.0679), indicating an absence of NOS-dependent stiffening in the female myocardium.

Systolic ventricular pressure and indices of dP/dt were unchanged by knockdown in both male and female hearts, indicating preserved systolic function (Figure 3F and 3G). In *Cavin-1*^ww^ male hearts, L-NAME treatment elicited a negative lusitropic effect, evidenced by a significant reduction in dP/dt_min_ (*p*_interaction_=0.0395; Figure 3G). In contrast, L-NAME did not alter functional indices in *Cavin-1*^ff^ males or in female hearts of either genotype (Figure 3G and 3H).

Coronary microvascular function was assessed from reactive hyperaemia responses following 15- and 30-sec global ischaemia. Hyperaemia was preserved in both male and female hearts irrespective of cavin-1 status (Figure 3I-L). Inhibition of NOS with L-NAME, however, elicited a sex-specific vasoconstrictive response, reducing AUC selectively in males (Figure 3I and 3K; Males: *p*_treatment_(15s)=0.0244, *p*_treatment_(30s)=0.0183; Females: *p*_treatment_(15s)=0.6082, *p*_treatment_(30s)=0.2105). This implies preserved microvascular function with cardiomyocyte caveolae deficiency.

### Disruption of cavin-1-dependent caveolae impairs post-ischaemic performance in female hearts

Cardiac efflux of LDH was increased in male hearts irrespective of genotype following ischaemia (*p*_ischaemia_=0.0306; *p*_genotype_=0.8047), whereas *Cavin-1*^ff^ female hearts exhibited exaggerated LDH efflux compared to *Cavin-1*^ww^ females (*p*_interaction_=0.0134; Figure 4B), reflecting exacerbated reperfusion injury. Coronary flow was not impacted by *Cavin-1* genotype or ischaemia (p_interaction_=0.9573; Figure 4C).

**Figure 4:**
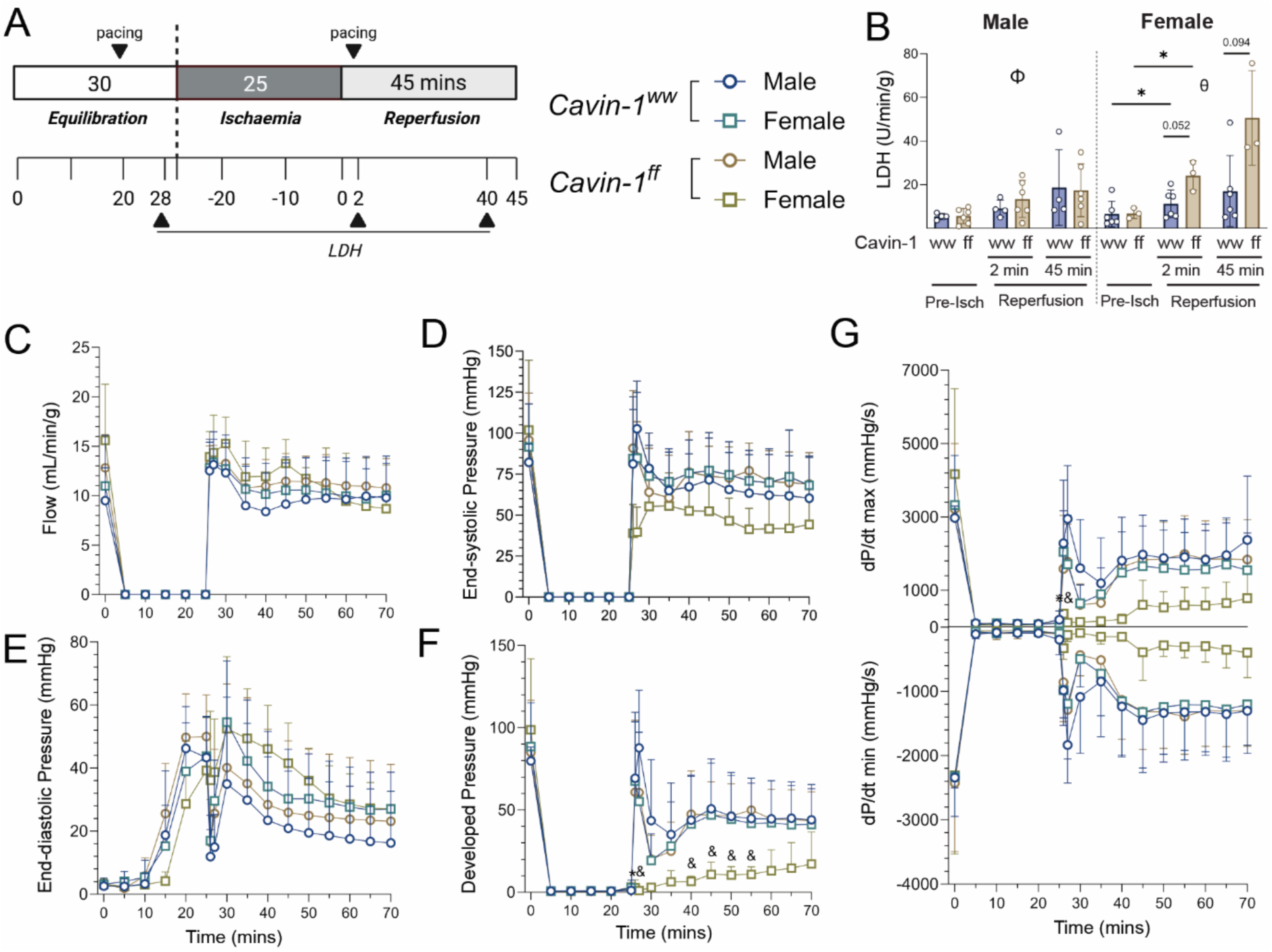
Ischaemic tolerance is impaired only in hearts of *Cavin-1*^ff^ females. (A) Schematic of the 25-minute Langendorff ischaemia-reperfusion protocol, indicating pacing and effluent collection timepoints for LDH analysis. (B) LDH efflux 2 minutes before ischaemia, 2 minutes into reperfusion, and after 45 minutes of reperfusion in male (left) and female (right) hearts. (C-G) Functional parameters across ischaemia and during reperfusion: (C) coronary flow rate, (D) end-systolic pressure, (E) end-diastolic pressure, (F) developed pressure, and (G) dP/dt_max_ and dP/dt_min_. Data are presented as mean + SD and were analysed by 2-way ANOVA with Tukey’s post hoc correction. *Cavin-1*^ww^ males: *n*=4, *Cavin-1*^ww^ females: *n*=7, *Cavin-1*^ff^ males: *n*=6, *Cavin-1*^ff^ females: *n*=3. *, *p*<0.05; θ, *p*_interaction_<0.05; Φ, *p*_ischaemia_<0.05. Multiple comparisons, &, *p*<0.05 vs *Cavin-1*^ff^ males; ⋇; *p*<0.05 vs *Cavin-1*^ww^ females.

*Cavin-1*^ff^ females exhibited impaired ischaemia tolerance, as indicated by reduced end-systolic pressure (*p*_interaction_=0.0.0156; Figure 4D), despite no corresponding effect observed for end-diastolic pressure (*p*_interaction_=0.0726; Figure 4E), eliciting a marked reduction in developed pressure throughout reperfusion (*p*_interaction_=0.0002; Figure 4F). Consequently, dP/dt_max_ (*p*_interaction_=0.0278), but not dP/dt_min_ (*p*_interaction_=0.0697), was impaired post-ischaemia (Figure 4G). *Cavin-1* knockdown had no effect on recovery from ischaemia in males.

## Discussion

We have undertaken a detailed characterisation of cardiomyocyte caveolae deficiency resulting from targeted cavin-1 disruption, revealing reduced ventricular compliance, increased wall thickness, reduced left ventricular systolic and developed pressures, impaired haemodynamic efficiency (stroke work/pressure-volume area), and increased heart rate. Importantly, we note a surprising systemic impact of cardiomyocyte caveolae deficiency, with reduced systolic blood pressure (despite elevated heart rate and preserved ejection) implying reduced peripheral resistance. We also provide the first comprehensive sex-stratified functional phenotyping of pathology arising from caveolar disruption following *Cavin-1* knockdown. In addition to the aforementioned effects in males and females, we document marked fibrosis in female but not male mice, whereas cardiac stiffness was selectively sensitive to NOS inhibition in male hearts. Depletion of cardiomyocyte cavin 1 also compromised ischemic tolerance in female, but not male, hearts.

### Systemic vs. cardiac functions of cardiomyocyte caveolae

We utilised multimodal assessment of cardiac function incorporating cardiac ultrasound and pressure volume analysis *in vivo* and *ex vivo*. Cardiomyocyte-specific cavin-1 deficiency caused a significant systemic effect, with reductions in systolic arterial blood pressure and total peripheral resistance despite fractional shortening comparable to wild-type mice. This was despite an elevated heart rate that was not a compensatory response to low blood pressure *in vivo*, as it was also present in isolated hearts. These systemic influences render interpretation of cardiac function *per se* more challenging. Profound changes in afterload (and heart rate) preclude direct comparison and interpretation of diastolic function *in vivo*, and highlight the utility of *ex vivo* analyses, where systemic influences are absent, heart rate can be normalised, and PV relationships examined at high resolution over a broad volume range. Nonetheless, decreased developed pressure and dP/dt *in vivo* indicates compromised ventricular performance, which was confirmed *ex vivo*. This is consistent with impaired electromechanical function (47), mimicking age-induced dysfunction (48, 49). Further, caveolin-3 and cavin-4, predominantly expressed in muscle tissue, have also been implicated in modulating excitation-contraction coupling and sarcolemmal integrity (32, 50–52), and their altered expression in this cohort may contribute to the nuanced dysfunction observed.

### Differing mechanisms and pathology in male versus female hearts

Sex divergent phenotypes included *Cavin-1*^ff^ female mice exhibiting reduced stroke volume *in vivo*, increased collagen deposition, and heightened susceptibility to ischemic injury compared with males. Further, males showed a greater diastolic dysfunction together with NO-dependencies of both ventricular stiffening and coronary function that were absent in females. These differences were not attributable to variation in AAV transduction efficiency, suggesting intrinsic sex-dependent modulation of caveolar signalling. Prior work supports sex differences in caveolin expression and function (53), with females slower to up-regulate NOS expression in response to mechanical stress, and sex differences in NOS isoform expression (54, 55). There is also evidence of differential hormonal regulation of myocardial remodelling pathways (56). Our findings extend these observations, identifying cavin-1 as a node through which sex-specific factors may influence cardiac structure and function.

Cardiac fibrosis is one recognised consequence of caveolar perturbations, documented in global knockout mouse models of *Cavin-1* (27), *Caveolin-1* (*57*), and *Caveolin-3* (13) but not *Cavin-2* (58). A key observation here is that cardiac fibrosis following *Cavin-1* knockdown is restricted to females, revealing a sex-dependent vulnerability. Of note, there is evidence that caveolae potentiate oestrogen receptor (ER) signal transduction (59), which contributes to cardioprotection and limits fibrotic remodelling (60, 61). Accordingly, the female-restricted fibrosis observed here may therefore reflect in part impaired ER-dependent antifibrotic signalling secondary to caveolar loss. As ER activity is also known to enhance basal NOS expression (62), this disruption may additionally alter the balance between ER-driven NOS induction and caveolae-mediated inhibition. However, direct assessment of the molecular precursors to fibrosis herein was constrained by a fixed timepoint analysis, though a trend towards female-specific elevation in baseline LDH efflux and troponin *ex vivo* aligns with known oestrogen-mediated effects on cellular integrity and stress responses (63, 64).

Reduction of cardiomyocyte caveolae may directly decrease compliance due to loss of the elastic membrane reserve these structures normally provide (21, 65). Caveolae contribute to effective sarcolemmal surface area, buffering mechanical load and supporting volume regulation. However estimates of the extent of this capacity vary; 12% in rat ventricular myocytes (65, 66), up to 27% (67), and by 56% in rabbit atrial myocytes (68). These higher estimates, however, seem inconsistent with the density of caveolae observed in the current study. The fibrotic response seen in females may thus reflect a detrimental adaptation to loss of sarcolemmal reserve leading to reduced membrane integrity and myocyte stiffening.

The sex-dependence of diastolic dysfunction here intersects with bimodal modulatory effects of NO on cardiac inotropy and lusitropy. End-diastolic pressure was disproportionately elevated at lower-range volumes in males, consistent with a functional impairment in ventricular compliance that was reversible with NOS inhibition. This pattern supports a role for NO-dependent modulation of sarcomeric Ca^2+^ in limiting early filling and raising EDP without gross structural change, an observation parallel to Kaakinen et al. (29). In contrast, females showed a NOS-independent shift of the EDPVR at high filling volumes, indicative of fixed structural impairment (likely increased myocardial stiffness from fibrosis) rather than a reversible biochemical dysregulation. These sex-dependent differences support divergent mechanisms of diastolic dysfunction with cardiomyocyte caveolae deficiency: NO-sensitive, functionally mediated compliance loss in males vs. fibrosis-driven, structural stiffening in females.

In *Cavin-1*^ww^ male hearts, NOS inhibition produced a marked reduction in contractility, reflected by decreased dP/dt indices, which underscores the role of NO in enhancing inotropy under normal physiological conditions. In contrast, *Cavin-1*^ff^ male hearts showed no systolic response to L-NAME, suggesting that NO levels in these hearts are elevated enough to decrease compliance, but not sufficiently high to confer an inotropic effect. Meanwhile, the absence of any functional effect of NOS inhibition in wild-type females implies sex-dependent redundancy or compensation in myocardial NO.

### Modulation of heart rate by caveolae

Elevations in heart rate were evident *in vivo*, persisted *ex vivo*, and were independent of NO signalling, indicating a chronic and intrinsic dysregulation unlikely to be driven by acute hormonal influences. Of relevance, caveolae act as signalling scaffolds that tightly regulate cardiac β-adrenergic signalling, together with sequestration of ion channels. Loss or disruption of caveolar components (for example *Caveolin-3* mutations or *Caveolin-1* deletion) potentiates the chronotropic response to β-adrenergic stimulation (69–71), consistent with receptor and effector sensitisation. Chronic sensitisation provides a plausible mechanism for tachycardia following cardiomyocyte-specific cavin-1 depletion, with loss of caveolar integrity increasing β-adrenergic drive or responsiveness, elevating heart rate in response to impaired haemodynamic performance. That said, persistence *ex vivo* (absent circulating catecholamines) also suggests shifts in the electrophysiology of atrial cells, where caveolae localise and modify ion channel function (including the pacemaker current) (72–74). Caveolar disruption (e.g. via cholesterol depletion) can increase heart rate by slowing deactivation kinetics of both HCN4 and *I*_f_, thereby accelerating diastolic depolarisation independently of cyclic nucleotide levels (72). Together, it seems cavin-1 deficiency dismantles signalling microdomains that typically constrain pacemaker channel activity thereby relieving parasympathetic restraint imposed by caveolae-mediated spatial organisation.

### Global versus cardiomyocyte-specific gene modulation

Prior studies of caveolar proteins have largely relied on germline knockout strategies. These approaches come with inherent limitations, including: 1) adaptation of the knockout colony to lifelong, multigenerational loss of function (75); 2) inability to discriminate between the protein role in developmental versus maturation versus adult heart function versus adaptation to stressors; and 3) inability to determine cell subtype-specific roles (76). Cavin-1 is ubiquitously expressed throughout the heart, most abundantly in endothelial cells (18), precluding detailed characterisation of myocyte-specific functions with global knockout. Global deletion of the muscle specific *Caveolin-3* is not lethal, however mice develop dilated cardiomyopathy with marked systolic dysfunction (13). Combined *Caveolin-1* and *Caveolin-3* deficiency exacerbates this phenotype and introduces vascular abnormalities (77), underscoring systemic contributions. The divergence from *Caveolin-3* knockout models here is noteworthy. Whereas *Caveolin-3* deficiency produces prominent ventricular dilatation (13), *Cavin-1* knockdown in cardiomyocytes yielded more subtle but multifaceted defects encompassing contractility, chronotropy, wall thickness and extracellular matrix remodelling (in females), suggesting that disruption of cardiomyocyte caveolae compromises mechanoelectrical signalling before gross chamber remodelling ensues. Further, caveolin/caveolae knockout models frequently exhibit endothelial dysfunction driven by altered NO bioavailability and excess ROS (78), whereas preserved reactive hyperaemia in the present model suggests endothelial function was maintained and was not contributing to cardiac dysfunction. This distinction suggests cavin-1 and caveolins, though interdependent for caveolae formation, may regulate partially non-overlapping signalling axes in cardiomyocytes.

The cardiomyocyte-specific role of caveolae is highlighted by several key observations. First, cardiac insufficiency and underload-induced hypotension were no longer obscured by impaired vasomotion or increased vascular stiffness (28), pointing to a heightened risk of tissue hypoperfusion. Second, the female-restricted fibrosis exposes a pivotal role for caveolae as signalling scaffolds governing sex-specific cardioprotective signalling. Third, lack of change in functional tolerance to ischaemia in male hearts, distinct from global *Cavin-1* KO models, emphasises the significance of microvascular integrity and whole-heart NO regulation in supporting recovery. In contrast, female hearts exhibited increased susceptibility to ischaemia-reperfusion injury, indicating a greater reliance on or influence of cardiomyocyte integrity and cardiac fibrosis.

Overall, divergence from global KO models, where endothelial and systemic compensations may obscure cardiac-specific effects, highlights the value of tissue-targeted approaches in dissecting caveolar protein function and represents a significant technical advance. Using AAV-inducible expression of iCre recombinase, we achieved a more physiologically relevant partial knockdown of *Cavin-1*, as opposed to comprehensive but biologically unrealistic complete gene knockouts. This offers a novel platform to investigate how cardiac dysfunction intersects with cardiomyocyte-specific caveolar depletion, as seen with ageing (79), and confirms a high dependence of caveolae formation on cavin-1. Moreover, limitations associated with germline deletions provoking developmental and generational adaptations or systemic confounders are highlighted by the phenotypic heterogeneity between seemingly identical cavin-1-null lines published recently (27, 29), whereas an incomplete, postnatal, cardiomyocyte-targeted reduction more closely resembles the gradual loss of caveolar integrity in acquired disease.

### Future directions

This novel model provides impetus for analogous strategies in other cardiac cell types. Fibroblast-specific knockdown could clarify how caveolae modulate extracellular matrix remodelling, whereas endothelial-specific models might reveal contributions to vascular–myocyte crosstalk and vessel integrity. The combination of floxed *Cavin-1* alleles with cell-specific and inducible Cre systems therefore provides a novel platform to explore to a more granular dissection of caveolar biology in cardiovascular health and disease. Integration with transcriptomic/proteomic approaches will further delineate the downstream signalling networks disrupted by cavin-1 loss.

From a translational perspective, our findings strengthen the rationale for targeting caveolae in cardiovascular disease. Strategies aimed at stabilising caveolar structures, enhancing cavin-1 expression, or modulating downstream signalling may hold promise for conditions characterised by impaired contractile reserve, diastolic dysfunction, maladaptive fibrosis, or heightened ischemic vulnerability. Observed sex differences further highlight the need to tailor such interventions according to sex-dependent biology.

### Conclusions

In summary, we demonstrate that disruption of cardiomyocyte caveolae via cell-specific cavin-1 knockdown impairs cardiac compliance, alters chronotropic regulation, and sex-dependently influences fibrosis and ischaemic tolerance in females. These findings refine our understanding of caveolar biology in the heart, highlight the importance of cell-type specificity, and establish a foundation for future mechanistic and translational studies. By enabling dissection of caveolar function at the cellular level, our work lays a scaffold for future targeted approaches to prevent and treat cardiovascular disease.

## Acknowledgements

MER and WGT are supported by the National Health and Medical Research Council (NHMRC; grant number 2021380). MER, JPH and WGT are supported by the Australian Research Council (ARC; grant number DP190102072 WGT, MER; DP200101152 MER, JPH, WGT). MER, JSR, JPH and WGT are supported by Diabetes Australia Grant (Y22G-REIM). RGP is supported by an Australian Research Council (ARC) Laureate Fellowship (FL210100107). We gratefully acknowledge the input of Dr Di Xia in generating the Cavin flox mouse.

