## Supplementary Data for "Cardiomyocyte caveolae govern myocardial function and sex-dependent regulation of ventricular compliance and resilience via cavin-1"

**SUPPLEMENTARY INFORMATION**

**Table S1: PCR primers for genotyping *Cavin-1* mouse line.**

| **Primers for genotyping** | **Sequence (5’-3’)** |
| --- | --- |
| Forward | TCTCACTAACAACCCCACAAGATTTG |
| Reverse | AGCCCTGAATCACCCGCTATATC |

**Table S2 Validated primer sequences and working concentrations for RT-qPCR.**

| Target | Primer | Sequence (5’-3’) | Volume (µL) in 10 µL reaction (10 µM) |
| --- | --- | --- | --- |
| *Cavin-1* | Forward | AGGCCGGCCAGATAAAGAAAC | 0.125 |
|  | Reverse | TTGCTGACGCTCAGTTTGGC |  |
| *eGFP* | Forward | AGATCCGCCACAACATCGAG | 0.125 |
|  | Reverse | TCTCGTTGGGGTCTTTGCTC |  |
| *iCre* | Forward | CAAGCTGGTGGAGAGATGGA | 0.25 |
|  | Reverse | AGAGTCATCCTTGGCACCAT |  |
| *Gapdh* | Forward | AGGTCGGTGTGAACGGATTTG | 0.5 |
|  | Reverse | TGTAGACCATGTAGTTGAGGT |  |
| *Cavin-2* | Forward | ACTGCCAGGTGAATGCTGTC | 0.25 |
|  | Reverse | ACGCTGTTCCATGTTGTGCT |  |
| *Cavin-3* | Forward | TGCTCTTCAAGGAGGAGACTG | 0.125 |
|  | Reverse | CATCCTCTGGCTGATCCTGGG |  |
| *Cavin-4* | Forward | ACCAAAGTCGAAACCAAGCAAGA | 0.25 |
|  | Reverse | GTTCTCGGGCAGGCTTCTGT |  |
| *Caveolin-1* | Forward | TAAATCACAGCCCAGGGAAACC | 0.5 |
|  | Reverse | CGGGTCTACGTATTTGCCCC |  |
| *Caveolin-3* | Forward | CACACGGATCTGGAAGCTCG | 0.5 |
|  | Reverse | GGCTCCGCAATCACGTCTTC |  |
| *Nos2* | Forward | CTGCCTTTCACCTTGGAGAC | 0.5 |
|  | Reverse | CGTTTCCTGGGGATGAGATA |  |
| *Nos3* | Forward | GCTTCAGGAAGTGGAGGCTG | 0.5 |
|  | Reverse | CTGCAGTCCCGAGCATCAA |  |
| *Nppa* | Forward | GCTTCGGGGGTAGGATTGACA | 0.25 |
|  | Reverse | GCTCAAGCAGAATCGACTGCC |  |
| *Nppb* | Forward | TTTGGGCACAAGATAGACCGGA | 0.125 |
|  | Reverse | CCAGGCAGAGTCAGAAACTGGA |  |
| *Col1a1* | Forward | GTACATCAGCCCGAACCCCA | 0.25 |
|  | Reverse | GGTGGACATTAGGCGCAGGA |  |
| *Col3a1* | Forward | TGGCACAGCAGTCCAACGTA | 0.25 |
|  | Reverse | GTTGGGGCAGTCTAGTGGCT |  |
| *α-Sma* | Forward | CCCAGGCATTGCTGACAGGAT | 0.25 |
|  | Reverse | TGCTGGAAGGTAGACAGCGAA |  |
| *Tgfβ1* | Forward | TGCTGACCCCCACTGATACG | 0.25 |
|  | Reverse | TGGGGCTGATCCCGTTGAT |  |
| *Pdgfrα* | Forward | CATCCAGGTTAAAGGTTGCTGACT | 0.25 |
|  | Reverse | GGGTAATAAGAGCTGGCAGGAG |  |

**Table S3: Sex-stratified whole-heart qPCR and western blotting expression data for Figure 1.** Data are presented as mean ± SD and were analysed by two-way ANOVA.

|  |  |  | **Cavin-1^ww^** | **Cavin-1^ff^** | ***p*_interaction_** | ***p*_genotype_** | ***p*_sex_** |
| --- | --- | --- | --- | --- | --- | --- | --- |
| **mRNA** | *n* | Male | 6 | 3 |  |  |  |
|  |  | Female | 3 | 4 |  |  |  |
|  | *Cavin-1* Heart | Male | 0.97 ± 0.12 | 0.82 ± 0.22 | 0.7886 | **0.0393** | 0.4061 |
|  |  | Female | 1.06 ± 0.17 | 0.86 ± 0.10 |  |  |  |
|  | *Cavin-1* CMs | Male | 0.91 ± 0.33 | 0.76 ± 0.34 | 0.1889 | **0.0373** | 0.8064 |
|  |  | Female | 1.09 ± 0.15 | 0.51 ± 0.08 |  |  |  |
|  | *eGFP* | Male | 40.81 ± 6.77 | 97.57 ± 29.58 | 0.9285 | **<0.0001** | 0.2222 |
|  |  | Female | 30.67 ± 17.75 | 85.87 ± 16.81 |  |  |  |
|  | *iCre* | Male | 50.74 ± 8.39 | 121.19 ± 35.02 | 0.706 | **<0.0001** | 0.1404 |
|  |  | Female | 38.99 ± 21.88 | 101.78 ± 17.11 |  |  |  |
|  | *Cavin-2* | Male | 1.02 ± 0.17 | 0.80 ± 0.17 | 0.9395 | **0.0214** | 0.7178 |
|  |  | Female | 0.99 ± 0.16 | 0.77 ± 0.15 |  |  |  |
|  | *Cavin-3* | Male | 0.99 ± 0.18 | 1.35 ± 0.46 | 0.6253 | **0.0487** | 0.8315 |
|  |  | Female | 1.08 ± 0.28 | 1.31 ± 0.17 |  |  |  |
|  | *Cavin-4* | Male | 1.06 ± 0.19 | 0.80 ± 0.21 | 0.8855 | **0.0382** | 0.3388 |
|  |  | Female | 0.94 ± 0.2 | 0.72 ± 0.2 |  |  |  |
|  | *Caveolin-1* | Male | 0.95 ± 0.12 | 1.00 ± 0.13 | 0.8898 | 0.6324 | 0.1278 |
|  |  | Female | 1.07 ± 0.12 | 1.10 ± 0.18 |  |  |  |
|  | *Caveolin-3* | Male | 0.93 ± 0.07 | 0.79 ± 0.12 | 0.8922 | **0.0035** | 0.5903 |
|  |  | Female | 0.96 ± 0.07 | 0.81 ± 0.09 |  |  |  |
|  | *NOS2* | Male | 0.98 ± 0.13 | 1.24 ± 0.16 | 0.5502 | **0.0416** | 0.9496 |
|  |  | Female | 1.03 ± 0.07 | 1.18 ± 0.27 |  |  |  |
|  | *NOS3* | Male | 0.96 ± 0.13 | 0.93 ± 0.06 | 0.7228 | 0.8157 | **0.0184** |
|  |  | Female | 1.06 ± 0.01 | 1.07 ± 0.06 |  |  |  |
| **Protein** | *n* | Male | 4 | 3 |  |  |  |
|  |  | Female | 3 | 4 |  |  |  |
|  | Cavin-1 | Male | 1.10 ± 0.19 | 0.77 ± 0.08 | 0.3958 | 0.1657 | 0.4855 |
|  |  | Female | 0.87 ± 0.25 | 0.79 ± 0.37 |  |  |  |
|  | Caveolin-1 | Male | 0.93 ± 0.07 | 0.67 ± 0.19 | 0.2733 | 0.6059 | 0.0541 |
|  |  | Female | 1.09 ± 0.42 | 1.19 ± 0.37 |  |  |  |
|  | Caveolin-3 | Male | 1.06 ± 0.13 | 0.62 ± 0.42 | 0.8621 | **0.0482** | 0.5902 |
|  |  | Female | 0.93 ± 0.39 | 0.55 ± 0.36 |  |  |  |
|  | iNOS | Male | 1.11 ± 0.18 | 1.09 ± 0.27 | 0.4606 | 0.5334 | 0.3319 |
|  |  | Female | 0.86 ± 0.33 | 1.06 ± 0.26 |  |  |  |
|  | eNOS | Male | 1.14 ± 0.34 | 1.80 ± 0.70 | 0.6983 | **0.0300** | 0.0879 |
|  |  | Female | 0.81 ± 0.20 | 1.29 ± 0.34 |  |  |  |
|  | p-eNOS/ eNOS | Male | 0.89 ± 0.41 | 0.57 ± 0.22 | 0.6752 | **0.0425** | 0.2959 |
|  |  | Female | 1.15 ± 0.40 | 0.68 ± 0.16 |  |  |  |

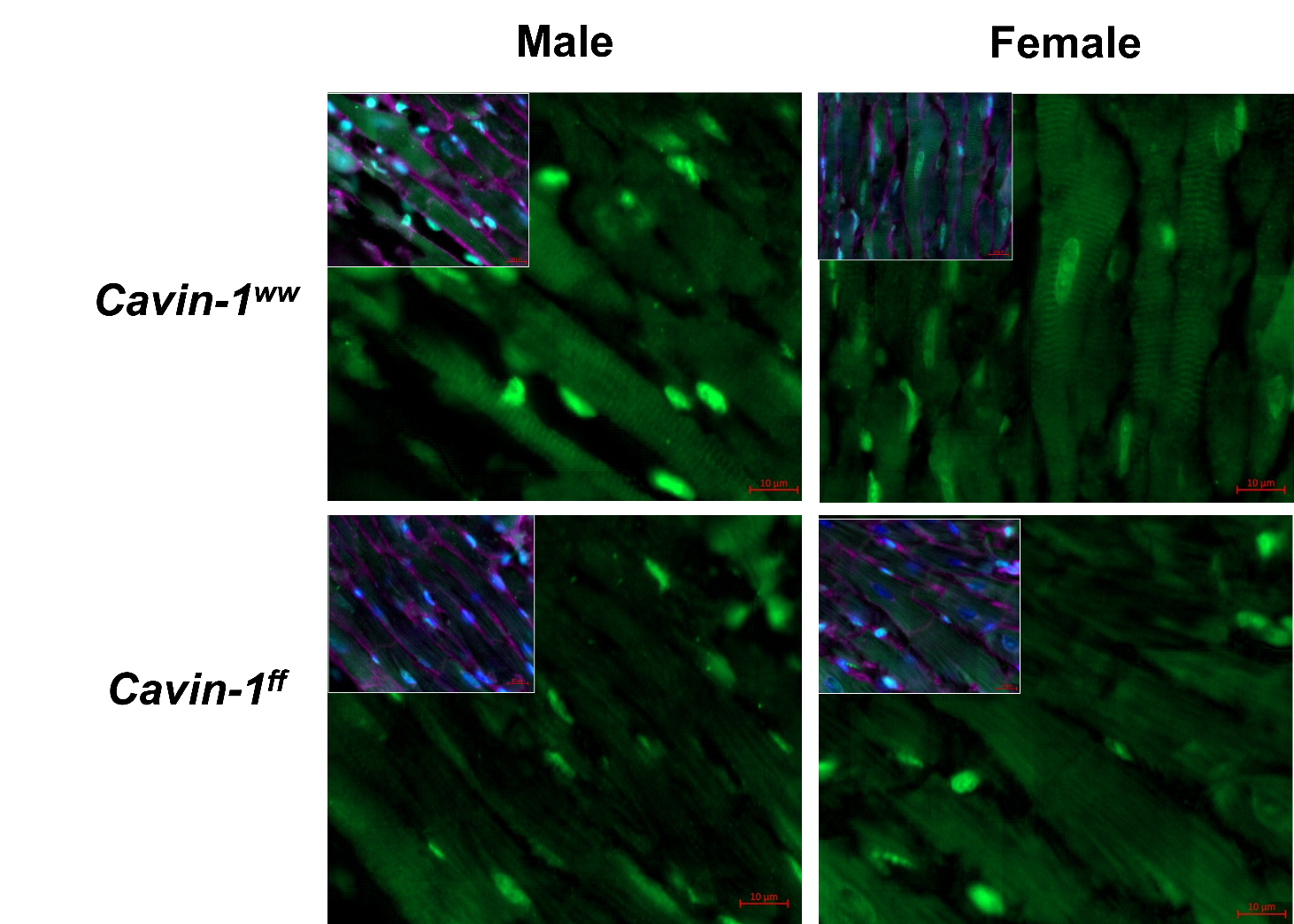

**eNOS WGA Hoescht**

**Figure S1: Immunofluorescent micrographs indicating eNOS mislocalisation in Cavin-1^ff^ hearts.** eNOS (green), wheat germ agglutinin (purple), and Hoescht (blue) staining in 5 μm paraffin mouse heart sections. Scale bars: 10 μm.

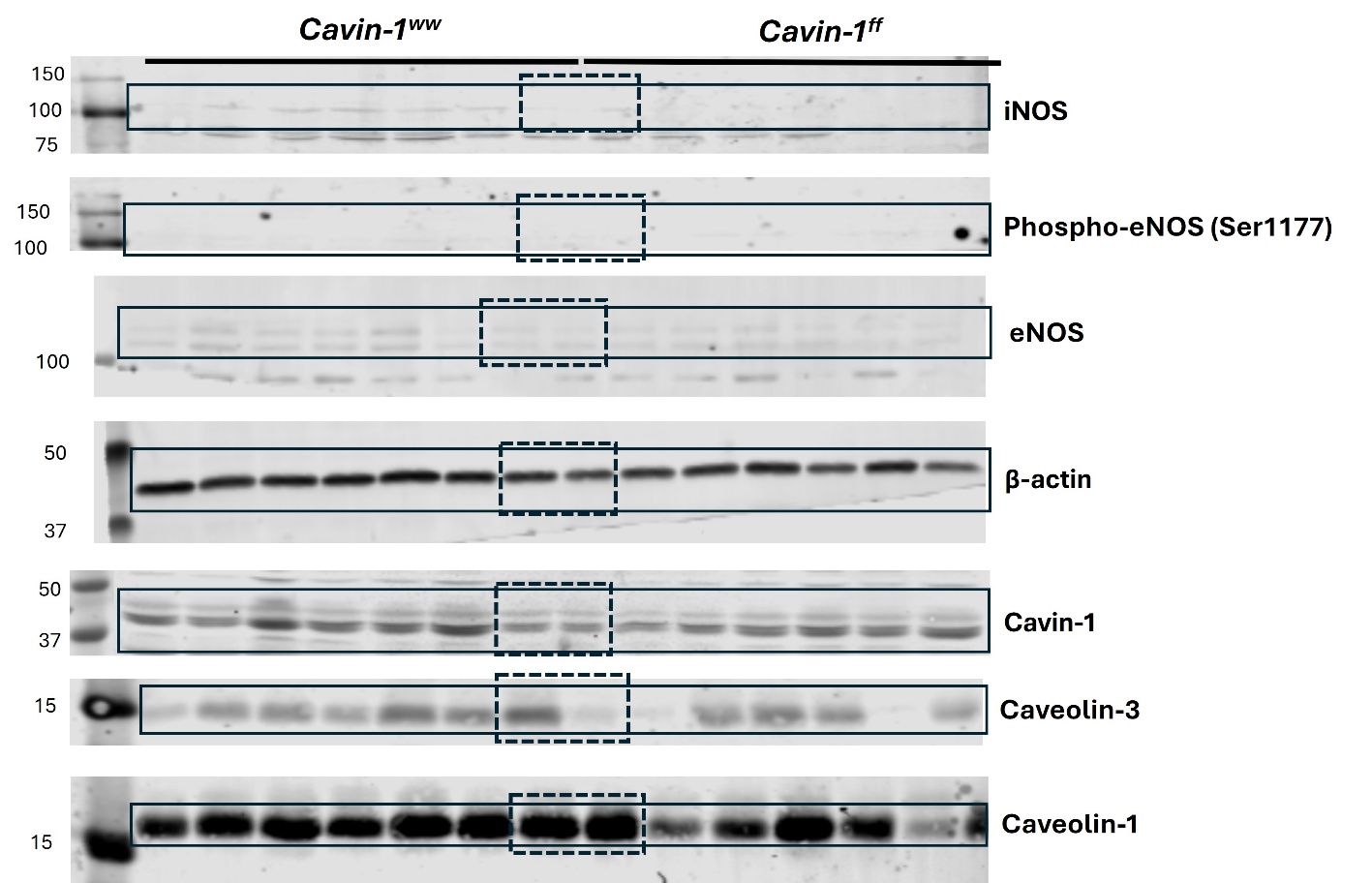

**Figure S2: Western immunoblots for Figure 1.** Protein expression is the average of two replicate immunoblots run concurrently. Membranes were sectioned according to molecular weight ladder and blotted independently.

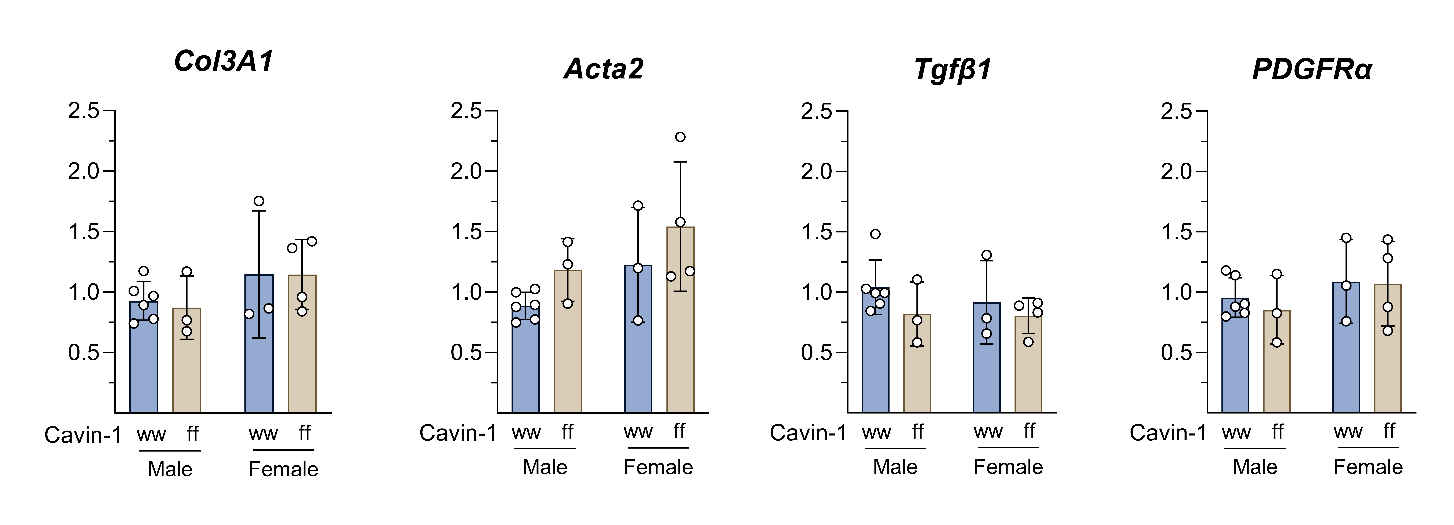

**Figure S3: mRNA expression of Col3a1, Acta2, TGFβ1, and PDGFRα.** Data shown as mean ± SD, and analysed by 2-way ANOVA.

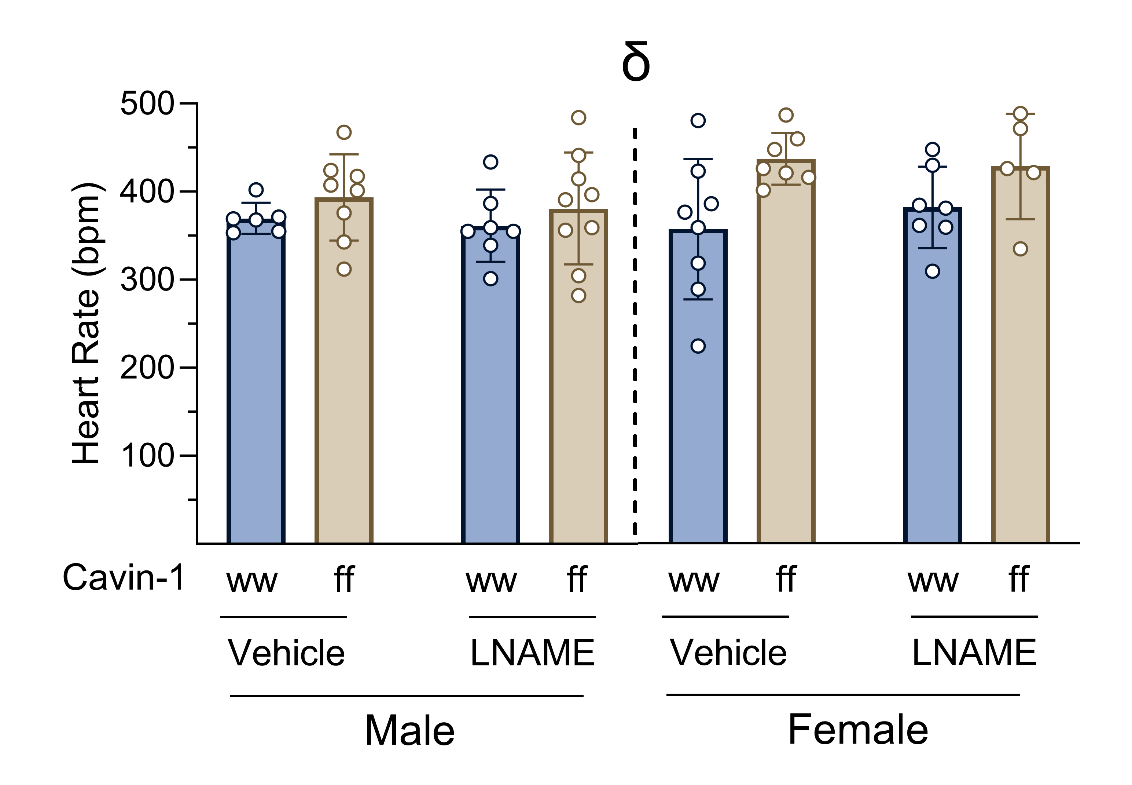

**Figure S4: Changes in heart rate ex vivo are NO-independent.** Basal heart rate shown before and after 100 µM L-NAME treatment in Langendorff perfused heart preparations. Analysed by 3-way ANOVA. p_treatment_=0.9353, **p_genotype_=0.0045**, p_sex_=0.0869, p_treatment x genotype_=0.5123, p_treatment x sex_=0.5161, p_genotype x sex_=0.1523, p_treatment x genotype x sex_=0.6115. Data shown as mean ± SD. Male: n_ww_=6, n_ww+L-NAME_=7, n_ff_=8, n_ff+L-NAME_=9; Female: n_ww_=8, n_ww+L-NAME_=7, n_ff_=7, n_ff+L-NAME_=5. δ: p_genotype_<0.05.

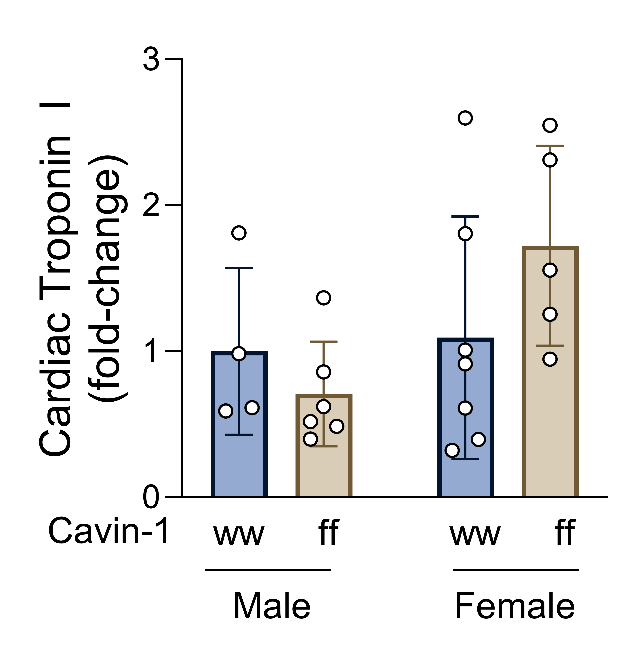

**Figure S5: Troponin release in isolated-perfused hearts.** Absorbance measured at 450 nm, normalised to flow and heart weight. Data presented as fold change vs. *Cavin-1^ww^* males, shown as mean ± SD, and analysed by 2-way ANOVA.
